# Faster amylin aggregation on fibrillar collagen hastens diabetic progression through β cell death and loss of function

**DOI:** 10.1101/2024.08.10.607320

**Authors:** Md Asrafuddoza Hazari, Gautam Kannan, Akash Kumar Jha, Musale Krushna Pavan, Subrata Dasgupta, Farhin Sultana, Soumya Ranjan Pujahari, Simran Singh, Sarbajeet Dutta, Sai Prasad Pydi, Sankhadeep Dutta, Prasenjit Bhaumik, Hamim Zafar, Ashutosh Kumar, Shamik Sen

## Abstract

Amyloid deposition of the neuroendocrine peptide amylin in islet tissues is a hallmark of type 2 diabetes (T2DM), leading to β-cell toxicity through nutrient deprivation, membrane rupture and apoptosis. Though accumulation of toxic amylin aggregates in islet matrices is well documented, the role of the islet extracellular matrix in mediating amylin aggregation and its pathological consequences remains elusive. Here, we address this question by probing amylin interaction with collagen I (Col I)—whose expression in the islet tissue increases during diabetes progression. By combining multiple biophysical techniques, we show that hydrophobic, hydrophilic & cation-π interactions regulate amylin binding to Col I, with fibrillar collagen driving faster amylin aggregation. Amylin-entangled Col I matrices containing high amounts of amylin induce death and loss of function of INS1E β-cells. Together, our results illustrate how amylin incorporation in islet matrices through amylin-Col interactions drives T2DM progression by impacting β-cell viability and insulin secretion.

## Introduction

Amyloid fibrils, which have become notably associated with diseases such as Alzheimer’s, Parkinson’s, Huntington’s disease, and Diabetes, are formed by 36 different human proteins that have been linked to more than 50 diseases ^1^. The intrinsically disordered peptide hormone islet amyloid polypeptide (IAPP), also known as amylin, is composed of 37 amino acids ^2^. In 90% of type II diabetes cases, IAPP is found as amyloid aggregates surrounding β-cells in the pancreas ^3–5^. Extensive research has demonstrated a strong correlation between the aggregation of human IAPP (hIAPP) and the death of insulin-secreting pancreatic β-cells during the progression of type II diabetes ^6–8^. Further evidence for this link is provided by comparing human and mouse IAPP: mouse IAPP differs from its human counterpart by six residues, three of which are β-strand-breaking prolines. Consequently, mouse IAPP does not form amyloids ^9,10^. Furthermore, mice engineered to express human IAPP and fed a high-fat diet can be induced to develop islet amyloid and type II diabetes ^11,12^. Studies using cell models and transgenic rodents have found that hIAPP fibrils are associated with apoptosis, β-cell loss, and severity of type II diabetes ^13–18^. Conversely, other studies suggest that the process of hIAPP fibril formation, rather than the fibrils themselves, is the primary source of toxicity ^19,20^.

It has been suggested that amyloids in different tissues may form, at least in part, due to interactions with major components of the extracellular matrix (ECM) specific to those tissues. These include ECM components like collagens I, II, IV, XXVA1, XVIII ^21–26^ and glycosaminoglycans (GAGs) ^27–29^. The binding affinities of amyloidogenic proteins/peptides to these collagens have been shown to be in the μM− mM range ^30^, with a preference for fibrillar collagens ^31^. The pathological significance of ECM-amyloid associations is supported by various studies. For instance, images of ex-vivo dialysis-related amyloidosis (DRA) plaques reveal amyloid covering the surface of collagen I fibrils ^24^. Alzheimer’s disease pathology is promoted through the consistent co-aggregation of specific collagens with Aβ plaques, and the compaction of Aβ plaques is influenced by collagenous amyloid plaque components (CLACs) derived from the collagen type XXV a1 chain (COL25A1) ^25^. In type 2 diabetes, amylin deposition between β cells and the islet vascular ECM disrupts islet capillary morphology, causes toxicity to surrounding endocrine cells, and decreases islet capillary density ^32–34^. An electron microscopy (EM) study in hIAPP (’HIP’) TG rats also reported morphological abnormalities in the vasculature of amyloid-laden islets ^35^.

Recent kinetic studies have shown that ECM components, such as collagen ^24,29,36^ and glycosaminoglycans (GAGs) ^37–39^, as well as preformed fibril seeds and other cofactors ^40–44^, regulate the amyloid formation of amylin and other proteins. However, the atomic intricacies of how these components interact with and instigate amyloid formation of amylin have remained an unresolved question. Furthermore, the consequences of these ECM-entangled amyloids on cellular health remain to be elucidated. Investigating such queries necessitates in vitro modelling of the interaction between cells and amyloid-entangled ECM. The monomeric amylin measures approximately 4.6 × 1.6 × 1.6 nm, whereas the simplest triple helical unit of collagen I boasts significantly larger dimensions of 300 nm × 1.5 nm × 1.5 nm. Collagen I triple helices assemble into even larger, structured fibrils ranging from 10 to 500 nm in diameter and μm-scale in length. Consequently, collagen I presents a vast surface with numerous reactive groups for amylin interactions.

In this study, we address the importance of amylin-collagen I interactions in the progression of type II diabetes. Using single-cell RNAseq data from healthy, prediabetic, and diabetic subjects, we first establish a correlation between amyloid deposition of amylin in the ECM with diabetes progression and then validate the same in a diabetic mice model. Using multiple biophysical techniques, we establish a robust interaction between amylin and collagen I identify the binding interfaces of amylin for collagen I at physiological pH. Using aggregation kinetics experiments, we demonstrate that fibrillar collagen I accelerate amyloid formation, with the amyloid fibrils remaining associated with collagen I. We then establish the pathological implications of amylin-associated collagen by demonstrating its impact on loss of β-cell viability and functionality. Our study illustrates how fibrillar collagen I drive amylin aggregation and, subsequently, progression of diabetes through β cell death.

## Materials and methods

### scRNA-seq analysis

We collected pre-processed human pancreatic single-cell RNA-Seq data from the Human Pancreas Analysis Program ^45^ database (https://hpap.pmacs.upenn.edu), encompassing 155,801 cells from 46 donors with normal and T2D annotations. We applied the scDREAMER ^46^ integration method to this data in its semi-supervised setting to infer the batch-corrected cellular latent embeddings. For running scDREAMER, we used the provided cell type annotations as labels and treated unknown cell types as missing labels. The latent embeddings inferred by scDREAMER were utilized for UMAP visualization of the cells.

We categorized the cells into different biological conditions: Normal, Pre-Diabetic, and Diabetic, based on the HbA1c condition with the normal and T2D samples that were already annotated in the database. The criteria for categorization were as follows: Normal (HbA1c ≤ 5.6 mmol/L), Pre-Diabetic (5.7-6.4 mmol/L), and Diabetic (≥ 6.5 mmol/L) ^47–50^. We further analyzed the Beta cell subset, which consisted of 34,958 cells. Using the Dot Plot visualization tool, we compared the expression level differences of genes associated with β−cell functionality (FDX1, MAFA, NEUROD1 and PAX6), Cell proliferation (PITX2, CTNNB1, CCND2, and MKI67), processing (ABCC8, KCNJ11, and SLC30A8), and mitochondrial stress (EIF2AK3, ATF4, and, HSPA5) across the three different conditions.

### Pancreas isolation and fixation

Male and female C57BL/6NJ (JAX - Strain #:005304) cohort mice were fed a regular chow (RC), kept on a 12-hours light/dark cycle, and maintained at room temperature (23°C). Mice consumed a high-fat diet (Research diet: D12492 Rodent Diet With 60 kcal% Fat) for 16 weeks from 8 week of their age. Mice were euthanized for the isolation of whole pancreatic tissue and fixed for 48 hours in 4% paraformaldehyde.

### Tissue processing, H&E staining & Immunofluorescence

At first, the pancreatic tissue was placed overnight in a fresh 50 ml Falcon tube containing 30 mL of a 15 % sucrose solution. The next day, the organ was stored in a 30% sucrose solution for the whole night. The tissue was cryosectioned (5µm) and taken on microscopic slides coated with poly-l-lysin the following day.

H&E staining was performed according to standard procedure. The slides were put into 1x PBS for 3 minutes, followed by a gentle wash with tap water. The slides were then stained with Mayer’s Haematoxylin for 2 min, followed by washing with tap water for 4 min until the nuclei were stained blue. After placing in 95% alcohol, counterstaining was performed with an alcoholic Eosin solution for 2 min, followed by dehydrating through 2 changes of 95% EtOH and 3 changes of 100% EtOH. After placing in xylene solution for 1 min, the sections were mounted with mounting media, followed by visualization under a light microscope.

For staining, the slides were kept at 60°C for 10 min. After cooling at room temperature, the sections were washed in 1x PBS. Each tissue section was outlined with a hydrophobic barrier pen by placing the slides with the tissue facing up in a humidified chamber. The sections were then incubated with a sufficient volume of 8-9% BSA-blocking solution overnight. After that, primary antibodies were added to the slides and incubated in the humidified chamber at 4°C freezer overnight. Slides were washed three times with 1x PBS in 0.1% TritonX-100, followed by incubation with respective secondary antibodies at a dilution of 1:1000 for 3-4 hours at room temperature. Afterward, the slides were washed three times with 1x PBS in 0.1% TritonX-100. The nuclear staining was performed with Hoechst staining solution for 3 min, followed by washing three times in 1x PBS. Finally, the sections were mounted with mounting media, followed by sealing with nail polish on all edges, thereby, allowing slides to dry completely before imaging. Primary antibodies used were: mouse anti-Insulin (1:300) and rabbit anti-collagen (1:200).

#### Chemicals and Reagents

Details of chemicals and reagents used for the study are listed in Table 1.

**Table 1:** Catalog numbers of reagents, kits used in the study.

| Reagent type or resources | Designation | Source or reference | Identifiers | Additional information |
| --- | --- | --- | --- | --- |
| Cell line | INS1E | gifted by Prof. Claes Wollheim and Prof. Pierre Maechler from the University Medical Centre, Geneva, Switzerland |  |  |
| Biological samples (Mus musculus) | Mice pancreatic Islet sections | IIT Kanpur, Institutional Animal Ethical Committee (IAEC) at IIT Kanpur |  |  |
| BrCn | Cyanogen Bromide | Sigma | 506-68-3 |  |
| HFIP | 1,1,1,3,3,3-Hexafluoro-2-propanol | Sigma | 920-66-1 |  |
| Collagen I | Type I Collagen | Corning® | 354249 |  |
| FITC | Fluorescein 5(6)-isothiocyanate | Sigma | 27072-45-3 | 3:1 (molar ratio) |
| Alexa fluor 546 dye | Alexa Fluor™ 546 NHS Ester (Succinimidyl Ester) | Thermo | A20002 | 3:1 (molar ratio) |
| Poly-L-lysine | Chemical | Sigma | P4707 |  |
| Glutaraldehyde | Chemical | Sigma | G7651 |  |
| Thioflavin T | Dye | Sigma | T3516 |  |
| RPMI-1640 | Cell culture media (CCM) | Gibco | 31800-022 |  |
| No glucose RPMI-1640 | CCM | HiMedia | AT150 |  |
| Fetal bovine serum (FBS) | CCM supplement | Gibco | 10270-106 |  |
| HEPES | CCM supplement | Gibco | 15630080 |  |
| sodium pyruvate | CCM supplement | Gibco | 11360039 |  |
| 2-mercaptoethanol | CCM supplement | Gibco | 21985023 |  |
| Paraformaldehyde | Fixation chemical | Sigma | 30525-89-4 |  |
| DPBS | Phosphate buffer saline | Gibco | 21600010 |  |
| Anti-Caspase 3 | Primary antibody | Invitrogen | 700182 | 1:300 |
| Anti-Insulin | Primary antibody | SCBT | SC377071 | 1:300 |
| Anti-RIPK3 | Primary antibody | Abclonal | A5431 | 1:500 |
| Anti-MLKL | Primary antibody | Abclonal | A5579 | 1:500 |
| Alexa-fluor 488 anti-mouse IgG | Secondary Antibody | Invitrogen | A-32723 | 1:500 |
| Alexa-fluor 555 anti-rabbit IgG | Secondary Antibody | Invitrogen | A-21422 | 1:500 |
| Hoechst 33342 | Dye | Merck | B2261 | 1:5000 |
| Alexa Fluor 488 Wheat Germ Agglutinin (WGA) | Dye | Invitrogen | W11261 | 1:1000 |
| Alexa Fluor 647 Wheat Germ Agglutinin (WGA) | Dye | Invitrogen | W11261/W32466 | 1:500 |
| Alexa Fluor™ 555 Phalloidin | Dye | Invitrogen | A34055 | 1:500 |
| Eukitt quick-hardening mounting medium | Mounting medium | Sigma | 03989 |  |
| CalceinAM | Dye | Invitrogen | C1430 |  |
| Propidium Iodide (PI) | Dye | Sigma | P4170 |  |
| 2-Deoxy-2-[(7-nitro-2,1,3-benzoxadiazol-4-yl) amino]-D-glucose (2NBDG) | Fluorescently tagged glucose | Sigma | 186689-07-6 |  |
| DCFDA (2',7'-Dichlorofluorescein diacetate) | Dye | Sigma | 4091-99-0 |  |
| TRIzol™ | Reagent for RNA, DNA, protein isolation | Invitrogen | 15596026 |  |
| cDNA Reverse Transcription Kit | Kit | Applied Biosystems | 4368814 |  |
| PowerUp™ SYBR Green Master Mix | PCR master mix | Applied Biosystems | A25742 |  |

#### Amylin expression and purification

The pGEV1-Amylin1-37 construct, which encodes the hexahistidine-tagged Amylin1-37 fused with GB1 (Protein G, B1 domain) and flanked by methionine at both the N- and C-termini, was transformed into *E. coli* BL21 (DE3) cells. Expression was carried out in 2xYT medium, and subsequent purification was performed according to previous protocol ^51^. In summary, cyanogen bromide (CNBr) was used to cleave the GB1, 6X-His tag, and additional methionine residues from both termini of the recombinant Amylin1-37 following protein elution, as previously described^52^. Post-cleavage, the resulting sample underwent ultracentrifugation (Beckman Coulter OptimaTM MAX-XP, USA) at 88,300 g for 55 minutes at 10°C. The resulting pellet was subsequently subjected to 4-5 washes with phosphate buffer and processed for monomer formation. Monomers were obtained by dissolving the pellet in HFIP (at 25 °C) and ultracentrifuging at 60,000 rpm for 3 hours. Following ultracentrifugation, the supernatant was collected and lyophilized overnight. The lyophilized protein was then reconstituted in HBS (pH 6.5) and further processed for subsequent interaction, fibril formation, and other experimental studies.

#### Molecular dissection of Collagen I/Amylin interactions through SPR

The molecular basis of interactions between Collagen I (Col I) and Amylin was probed using Surface plasmon resonance (SPR), FITC-based fluorescence assay and NMR. For SPR experiments, Collagen I (Col I) (Cat# 354249, Corning) monomers were immobilized onto Biacore CM5 sensor chips (GE Healthcare). Specifically, 100 µg/ml of Col solution prepared in 50 mM sodium acetate buffer at pH 4.5 was injected onto the chips at a rate of 5 µl/min for a duration of 773 seconds, resulting in a response unit (RU) of 39262. Next, varying dilutions of Amylin (20, 40, 60, and 80 µM) in HBS buffer containing 150 mM NaCl at pH 6.2 were subsequently injected over the immobilized Col on the chip at a flow rate of 20 µl/min for a duration of 90 sec at 10^0^ C. The dissociation phase was initiated at a flow rate of 20 µl/min, and the resulting signal recorded for 600 seconds. The chips were regenerated using 10 mM NaOH solution. To ensure accuracy, the data were subjected to reference subtraction to compensate for any differences in bulk refractive index. Furthermore, a consistent buffer system was employed for all proteins to avoid any changes in bulk viscosity. The kinetics data for the protein interactions were analyzed using a 1:1 two-state kinetic model within the Biacore T200 Software. This analysis allowed for the determination of Kon (association rate constant), Koff (dissociation rate constant), and KD (equilibrium dissociation constant) values, providing a better understanding of the protein-protein interactions. The fitted curves resulting from this analysis were plotted using Sigmaplot v.11.0 software.

#### Labelling of Amylin and Collagen I

We utilized FITC/Alexa Fluor 546 NHS Ester amine-reactive probes (Thermo Fisher Scientific, A20002) to efficiently label proteins at primary amines (located at the N-terminus of the polypeptide chain and in the side-chain of lysine (Lys)). The LMW protein solution was prepared using the manufacturer’s recommended non-amine-containing conjugation buffers (PBS, pH 9.5), and FITC & Alexa Fluor 546 labelling of LMW AMYLIN & Col respectively, was done following the manufacturer’s instructions (Thermo Fisher, Molecular Probes, USA). To label the proteins, we added three molar excess of amine-reactive FITC & Alexa Fluor 546 to the LMW protein solutions and incubated the protein-dye mixture for 2 hours at room temperature followed by 6 hours at 4 °C with continuous stirring. After that, we removed the free unreacted dye by extensive dialysis (∼ 36-40 hours, at 4 °C) in HBS (for AMYLIN, at pH 6.8) & 20 mM Acetic acid (for Col, at pH 4) buffer, using a mini dialysis unit (3.5 kDa MWCO, Millipore) with buffer exchange after every 4 hours. We measured the concentration and degree of labelling of the labelled protein using the manufacturer’s protocol (Molecular Probes, USA) and the equation (1) and (2). These were then used as FITC labelled protein stocks for further experiments.

A_protein_ = A_280_ – A_max_(CF) where A280 is the absorbance of the protein at 280 nm, A_max_ is the absorbance of the dye at its excitation maximum (λ^max^) and CF is the correction factor; CF for FITC & Col are 0.35 & 0.12 as listed in the manufacturer’s protocol.

Degree of labelling (D. O. L.): DOL = A_max_ x MW/ ([protein] x ε_dye_)

where MW = the molecular weight of the protein, A_max_ is the absorbance of the dye at its excitation maximum (λ^max^), ε_dye_= the extinction coefficient of the dye at its absorbance maximum, and the protein concentration in mg/ml ^53^.

#### FITC based fluorescence intensity assay: Amylin-Col interaction profile

For interaction studies, we have used FITC labelled Amylin dissolved in HBS buffer (pH 6.2) to monitor change in fluorescence intensity upon the addition of Col I at 10 °C. We kept the Amylin concentration consistent (i.e. 5 µM) across all conditions and varied Collagen I concentrations (0 – 0.2mg/ml). To monitor fluorescence intensity change we have used spectrofluorometer (Fluoromax 4) to set an excitation wavelength of 490 nm (excitation wavelength of FITC) and a spectral emission range (500 – 600 nm).

#### Atomic Force Microscopy (AFM) for measuring the strength of interaction

For the assessment of Amylin-Collagen I binding using AFM, spherical probes (67 kHz, Novascan) with a diameter of 4.5 μm were initially coated with a 10 μg/ml solution of poly-L-lysine (Sigma, Cat # P4707) for 20 minutes, followed by treatment with 0.5% glutaraldehyde (Sigma, Cat # G7651) for 20 minutes ^54^. Subsequently, the probes were immersed in a 0.4 mg/ml (100 µM) solution of Amylin peptide for 30 minutes, rinsed 3–4 times with filtered autoclaved distilled H_2_O, and then subjected to vacuum desiccation for 30 minutes. Following tip calibration using the thermal noise method, investigations into Amylin-Collagen I binding were conducted. During indentation, the probe was maintained in contact with the Collagen I-coated surface for 5 seconds to facilitate the formation of Amylin-Collagen I bonds, after which it was retracted at a tip speed of 3–4 μm/sec. The analysis of maximum adhesion force was carried out using Igor Pro 6.22 A software ^55^.

### Thioflavin T (ThT) assay: Amylin & Amylin-Collagen I aggregation kinetics

Amyloid fibril formation was assessed through ThT fluorescence assays involving Amylin, either with or without the presence of Collagen I fibrils. To prepare the Amylin solution, lyophilized powder of purified recombinant Amylin was dissolved in 500 μL of HBS buffer at pH 6.5, resulting in a stock solution of 1.8 mg/mL (450 μM). This stock solution was further diluted to a concentration of 100 μM. Various concentrations of Collagen I fibrils were separately prepared by incubating Collagen I (at concentrations of 1.2 mg/mL, 0.5 mg/mL, or 0.1 mg/mL) in 50 μL of HBS at pH 7.4, maintained at 37°C for 1 hour in distinct wells of a 96-well plate to allow for gelation. Subsequently, Amylin was introduced into the wells, either with or without Collagen I, at a concentration of 100 μM. This process was repeated, generating three to six samples for each experimental condition within the 96-well plate. To monitor the fluorescence changes, ThT was added to each sample to reach a final concentration of 10 μM. The ThT fluorescence levels were tracked over a 24-hour period at 37 °C, with agitation at 180 rpm, using a synergy-H1 microplate reader from Biotek. The excitation wavelength was set at 450 nm, and the emission was measured at 482 nm. The lag time (t_lag_) was calculated utilizing the following equation, as reported in previous studies ^56^:

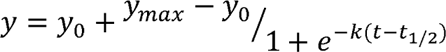

where:

- y represents the ThT fluorescence at a specific time point,
- *ymax* represents the maximum ThT fluorescence,
- y_O_ represents the ThT fluorescence at the initial time point (t_0_),
- The lag time (t_lag_) is defined as *t_lag_ = t_l/2_ - 2/k*

### Nuclear Magnetic Resonance (NMR) Experiments

Solid-state NMR experiments were carried out on a 3.2 mm HXY double resonance probe. The 1D 13C Cross-polarization magic angle spinning (CP-MAS) (36) experiment on Collagen I vs Amylin incubated Collagen I, was performed with an optimized cross-polarization (CP) contact time of ∼1.05 ms and a ramp of 100–50% on the spin-lock radio frequency (RF) power with a spinning frequency of 10kHz and a sample temperature of 10°C. A typical 1H 90° pulse of *ms RF of power 50 kHz was used. During 13C acquisition of up to 36 ms, the ^1^H RF power of Swept-frequency two-pulse phase modulation (SW_f_-TPPM)^57^ sequences with linear sweep profile for heteronuclear decoupling in solid-state NMR^57^. the decoupling scheme was set to 50 kHz. The recycle delay was kept for 2 s with an experimental time of 17.5 h with several scans of 30720.

For all liquid state NMR experiments, we prepared a solution of purified recombinant [U−^15^N]-labeled human islet amyloid polypeptide (Amylin) by diluting it to a final concentration of 100 μM in a buffer solution of HBS (pH 7), 10% v/v deuterium oxide (D_2_O). The concentration of Collagen I following dialysis was determined using a Nanodrop spectrophotometer. All NMR experiments were conducted at a temperature of 10 °C. The data acquisition took place using a 750 MHz Bruker AVIII NMR spectrometer. We obtained ^1^H−^15^N HSQC spectra for [U−^15^N]-labeled human islet amyloid polypeptide (Amylin) at various Collagen I concentrations (0, 0.2, 0.4, and 0.8 mg/mL) within an HBS buffer at pH 6.5. To assess the impact of Collagen I on the spectra, we calculated the intensity ratio for each cross-peak in the ^1^ H−^15^N HSQC spectrum of Amylin when Collagen I was present, relative to the intensity of the same cross-peak when Collagen I was absent. These intensities were determined using Bruker Topspin software, and errors were calculated from the experimental replicates ^58^.

#### Cell Culture Experiments

INS-1 cells (gifted by Prof. Claes Wollheim and Prof. Pierre Maechler from the University Medical Centre, Geneva, Switzerland) were cultured between passages 56 and 65. These cells were maintained in monolayer cultures under a humidified atmosphere containing 5% CO2 at 37°C. The culture medium consisted of RPMI-1640 (Gibco, Cat# 31800-022) supplemented with 10% fetal bovine serum (Gibco, Ref# 10270-106), 10 mM HEPES (Gibco, Cat# 15630080), 2 mM L-glutamine, 1 mM sodium pyruvate (Gibco, Cat# 11360039), and 0.05 mM 2-mercaptoethanol (Gibco, Cat# 21985023).

### Docking and MD simulations

The structural data for amylin monomer (PDB ID: 5MGQ)^103^ and amylin fibrils (PDB ID: 6VW2)^61^ were obtained from the RCSB Protein Data Bank. The collagen molecule structure was generated using the ColBuilder computational program^59^. Subsequently, each of these protein structures underwent energy minimization using the GROMOS96 force field in the Swiss-PDBViewer program, involving 3000 steps of conjugate gradient optimization followed by 2000 steps of steepest descent. The docking process involved the placement of the amylin monomer/fibril into the collagen monomer/fibril using the LightDock server. LightDock utilized the Glowworm Swarm Optimization algorithm to calculate protein-protein interactions.

The molecular dynamics simulations of protein-protein complexes were conducted using GROMACS v2022.2, employing the AMBER99SB-ILDN force field. Each complex was solvated, and the appropriate number of Na+ and Cl-ions were added for system neutralization. Energy minimization was performed using the steepest descent method with 50,000 steps and a tolerance of 1000 kJ mol-1 nm-1. Equilibration protocols involved 5 ns simulations under both constant particle number, volume, and temperature (NVT) conditions using a modified Berendsen thermostat with velocity rescaling at 310K and a time step of 0.1 ps, as well as constant particle number, pressure, and temperature (NPT) conditions utilizing Parrinello−Rahman pressure coupling at 1 bar with a compressibility of 4.5 × 10−5 bar−1 and a 2 ps time constant. The Nosé−Hoover thermostat was applied during NPT equilibration and the final production run to ensure the correct kinetic ensemble. Bond lengths were constrained using the Linear Constraint Solver (LINCS) algorithm. Long-range interactions were handled using the particle-mesh Ewald (PME) method with a 1.2 nm cutoff and a Fourier spacing of 0.16 nm. Subsequently, the equilibrated systems underwent a 100 ns final production run, with trajectory snapshots saved every 10 ps for subsequent data analysis.

#### Immunofluorescence microscopy

For immunostaining, cell fixation was performed after 48 hours of culture using 4% paraformaldehyde (PFA) in 1x phosphate-buffered saline (PBS) for 20–25 minutes. Subsequently, cells underwent three washes with 1x PBS to eliminate residual paraformaldehyde, followed by permeabilization with 0.1% Triton-X 100 (Sigma, Cat # 93443) in 1x PBS for 3 minutes. Blocking was carried out with 5% BSA (Sigma, Cat # A2058) for 2 hours at room temperature (RT) prior to overnight incubation at 4 °C with one of the following primary antibodies: Caspase 3 (Invitrogen, Cat# 700182), Insulin (SCBT, Cat# SC377071), RIPK3 (Abclonal, Cat# A5431), MLKL (Abclonal, Cat# A5579). The subsequent day involved three washes with 1x PBS, followed by incubation with one or more of the following secondary antibodies at room temperature for a minimum of 1–2 hours: Alexa-fluor 488 anti-mouse IgG (Invitrogen, Cat # A-32723), Alexa-fluor 555 anti-rabbit IgG (Invitrogen, Cat# A-21422). Nuclei and membrane were stained with Hoechst 33342 (Cat# B2261, Merck) (1:5000) and Alexa Fluor 488/ Alexa Fluor 647 Wheat Germ Agglutinin (WGA) (Invitrogen, Cat# W11261/ Cat# W32466) respectively for 15 minutes at RT. For F-actin staining post-secondary antibody incubation, Alexa Fluor 555-conjugated phalloidin (Thermo Fisher Scientific, Cat# A12380) was applied to the coverslips for 2 hours at room temperature in the dark. Following washes, cells were mounted on glass slides using Eukitt quick-hardening mounting medium (Sigma, Cat # 03989). Imaging at 63x magnification was conducted with an immunofluorescence microscope (Zeiss, LSM 780) for probing Caspase 3, RIPK3, MLKL, and Insulin. Image processing and analysis were carried out using Fiji ImageJ software ^60^.

#### Live dead assay

For live dead assay, 60,000 INS1E Cells were seeded on Collagen I hydrogel, Collagen I hydrogel incubated with Amylin (10 or 100 µM), and only Amylin-coated coverslips in a 24 well plate. After 48 hours of incubation, Cells were washed with PBS twice, and incubated with (4µg/ml) CalceinAM (Invitrogen, Cat# C1430) and (67 ug/ml) Propidium Iodide (PI) (Sigma, Cat# P4170) in plane RPMI 1640 for 30 minutes then washed with PBS. Subsequently, Cells were counter-stained with Hoechst 33342 (Cat# B2261, Merck) for 5 min in plane RPMI 1640 and washed with PBS & incubated in complete RPMI 1640 (Gibco, Cat# 31800-022) for imaging under an Olympus IX83 fluorescence microscope. Images were processed in ImageJ software for Live/dead cell profiling.

#### Glucose uptake assay

INS1E cells were incubated on various conditions (Collagen I/Collagen I+Amylin/ Only Amylin) for 48 hrs. Afterwards, Cells were washed with PBS and incubated with 100 µM 2-Deoxy-2-[(7-nitro-2,1,3-benzoxadiazol-4-yl) amino]-D-glucose (2NBDG) (Sigma Cat# 186689-07-6) containing RPMI1640 (HiMedia Cat# AT150) for 1 hr followed by wash with PBS and fixing in 4% PFA. Samples were further processed and stained with Hoechst 33342 & Alexa fluor 647-WGA (15 min., at RT) for fluorescence imaging under Olympus IX83 microscope. Images were processed & analysed by ImageJ software.

#### ROS assay

The DCFDA (2′,7′-Dichlorofluorescin diacetate) (Sigma Cat# 4091-99-0) assay protocol involves the diffusion of DCFDA inside cells which is deacetylated by cellular esterases to convert it in a non-fluorescent compound. Oxidation of such compound by ROS results fluorescent 2’,7’-dichlorofluorescein (DCF), which is proportional to the ROS levels in the cell cytosol. INS-1E cells were plated on Collagen I/Collagen I+Amylin/Only Amylin having coverslips in a 24-well plate (Nest) at an initial density of 60,000 cells/well and were allowed to grow at 37°C for 48 hrs. post-incubation, Cells were washed with PBS twice and incubated with DCFH-DA containing plane RPMI1640 30 minutes at 37 °C. After incubation cells were washed with PBS twice and incubated with complete RPMI 1640 and imaged in live condition.

#### Cell proliferation assay

INS1E cells were incubated on various conditions (Collagen I/Collagen I+Amylin/ Only Amylin) for 48 hrs. Afterward, Cells were washed with PBS and fixed with 4% PFA. Samples were further processed by incubating rabbit monoclonal Anti-Ki67 antibody (Invitrogen Cat# MA5-14520) at 4 °C overnight. Cells were washed with PBS and incubated with Alexa-fluor 555 anti-rabbit IgG at RT for 4 hours. Afterwards, Hoechst 33342 & Alexa fluor 647-WGA (15 min, at RT) staining were done and fluorescence images were taken under Olympus IX83 microscope. Images were processed & analysed by ImageJ software.

#### Quantitative PCR and gene expression array

Total RNA was isolated from INS1 cells employing TRIzol™ Reagent according to manufacturer’s protocol (Invitrogen, Cat # 15596026). Subsequently, 2 µg of total RNA underwent reverse transcription using the High-Capacity cDNA Reverse Transcription Kit (Applied Biosystems, Ref. # 4368814), followed by amplification using the PowerUpTM SYBR Green Master Mix (Applied Biosystems, Ref.# A25742) in a QuantStudio 5 instrument (Applied Biosystems). The analysis of PCR data was executed utilizing the comparative Ct method, with GAPDH expression serving as the reference for normalizing gene expression data. A list of primers can be found in a table of supplementary data.

## Results

### Increased expression of Collagen I (Col I) and Amylin during diabetes progression correlates with loss of islet p cells

To understand how alterations in the islet microenvironment correlate with alterations in f3 cell abundance and function during diabetes progression, we analyzed single-cell RNA sequencing (scRNA-seq) data from the Human Pancreas Analysis Program (HPPA) database ^45,46^. From the HPPA database, we collected a pre-processed dataset consisting of 155,801 cells from 46 donors categorized into normal, pre-diabetic and diabetic groups based on HbA1c levels (see Methods). After filtering and categorizing the data, we identified 81,596 cells in the normal condition, 26,453 cells in the pre-diabetic condition, and 47,752 cells in the diabetic condition across all cell types. We performed integration of the cells across donors using scDREAMER and identified 10 cell types: acinar cells, a cells, f3 cells, o cells, ductal cells, endothelial cells, E cells, immune cells, y cells and stellate mesenchymal.

We then focused on the f3 cell population consisting of 21,666, 6,707, and 6,585 cells from normal, pre-diabetic, and diabetic conditions, respectively, with a prominent reduction in the proportion of f3 cells [Figure 1A (highlighted region)]. Profiling of these cells revealed diabetes progression is associated with reduced f3 cell proliferation, increased mitochondrial stress, and an increase in expression of f3 cell functionality markers and amylin/insulin processing proteins (Figure 1B). Islet cells reside in islet tissue enriched in Col I secreted by islet endothelial cells and stellate cells raising the possibility that alterations in islet function is determined by alterations in the islet ECM ^35,62–67^. Consistent with this notion, increased expression of Col I (COL1A1 and COL1A2) in endothelial/stellate cells was correlated with increased insulin expression, increased amylin expression and increased insulin to amylin ratio under diabetic conditions (Figure 1C, Supp. Figure 1A, B). Moreover, a plot of Col I expression versus amylin expression revealed an increase in both the two from pre-diabetes to diabetes, suggesting alterations in ECM composition and amylin expression are positively correlated (Supp. Figure 1C).

**Figure 1:**
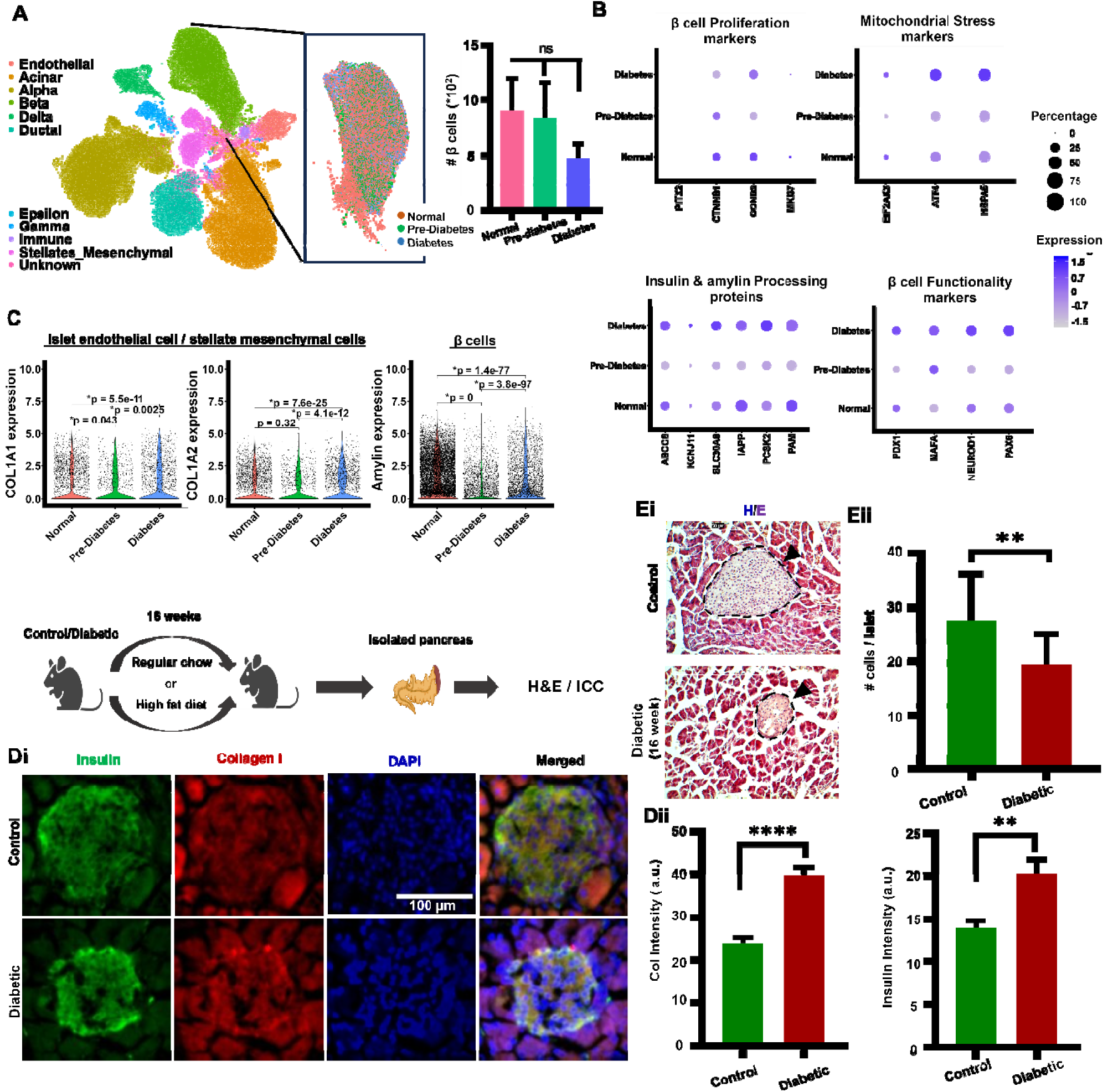
Alterations in Collagen/amylin expression impact islet organization and functionality during diabetic progression. (A) UMAP visualization of the Human Pancreas Analysis Program dataset. Cells are coloured based on the cell type, with the β cell subpopulation highlighted in which cells coloured based on diabetic condition. Also, the β cell distribution is shown across conditions. (B) Dot plot visualization of marker genes across different conditions, showing: (i) β cell Proliferation markers (ii) Mitochondrial stress markers, (iii) Insulin and amylin processing markers, and (iv) β cell Functionality markers. (C) The Expression levels of COLA1 & COL1A2 in Endothelial and Stellate Mesenchymal cells. Amylin expression levels in β cells. (Di) Experimental setup for treatment of mice to develop diabetes. Representative immunofluorescence images of islets from control and diabetic (16 week) mice in 5 μm-thick pancreas sections stained with DAPI (Nucleus), collagen I (red), insulin (green) and composite. Scale bar, 100 μm. (Dii) Quantification of mean collagen I and Insulin intensity across different conditions (≈ 15-22 Islets per condition, n=3) (Ei) Representative HE staining images of Islet from control and diabetic mice. (Eii) Quantification of cells per islet area from control and diabetic mice. (≈ 15-20 Islets per condition, n=3) (Data are presented as mean ± SEM. Statistical significance was assessed using One-Way ANOVA or nonparametric test (*p < 0.05,** p < 0.01,*** p < 0.001,**** p < 0.0001, ns: not significant).

To further probe the correlation between alterations in collagen expression and islet phenotype during diabetes progression in vivo, we performed H&E staining and immunofluorescence (IF) of pancreas isolated from control and 16-week diabetic mice. H&E images revealed a pronounced drop in the number of viable islet cells under diabetic conditions (Figure 1D). Further, quantification of pancreatic sections co-stained with Col I and insulin antibodies revealed increased expression of Col I in diabetic tissue and increased insulin expression in diabetic islets (Figure 1D). Collectively, these results suggest the temporal correlation of Col I and amylin expression is associated with the loss of islet f3 cells.

### Amylin interacts with both monomeric and fibrillar Col I

Given the increased expression of amylin and Col I under diabetic conditions, we hypothesized that their association may be a cause of f3 cell toxicity. To test our hypothesis, we first probed the nature of their association. Amylin is an intrinsically disordered protein. It contains two disordered regions, a shorter one at the N terminal (amino acid residues 1-5), and an extended disordered region at the C terminal (amino acid residues 17-37) interspersed by an a helical region (Figure 2A). Of the two disordered regions, the C terminal disordered region is most amyloidogenic (amino acid res. #25-30) followed by resideus14-19 located on the helix region as predicted by FoldAmyloid analysis. Col I is a triple helical peptide, composed of two a1 and one a2 chains, interwind to each other. These triple helices form fibrils by binding to each other. To probe the nature of interactions between amylin monomers and monomeric Col I (hereafter referred to as Col I), we monitored the change in fluorescence intensity upon Col I addition at increasing concentrations (0-0.2 mg/ml) to FITC-amylin (Figure 2Bi), with BSA serving as negative control. Drop in fluorescence intensity with increasing Col I concentration, but not with BSA, is indicative of amylin interaction with Col I with a characteristic K_d_ ≈ 1.5 µM. This association was further probed using SPR experiments, wherein amylin was flown over Col I-coated chips and the number of response units (RU) monitored. An increase in amylin concentration led to increase in RU with a two-state binding model yielding a value of K_d_ = 6.27 µM (Figure 2Bii). Consistent with this, confocal imaging of FITC-amylin incubated TRITC-Col I gels revealed prominent co-localization of FITC-amylin with TRITC-Col I (white arrows) (Fig. 2Ci, ii).

**Figure 2.**
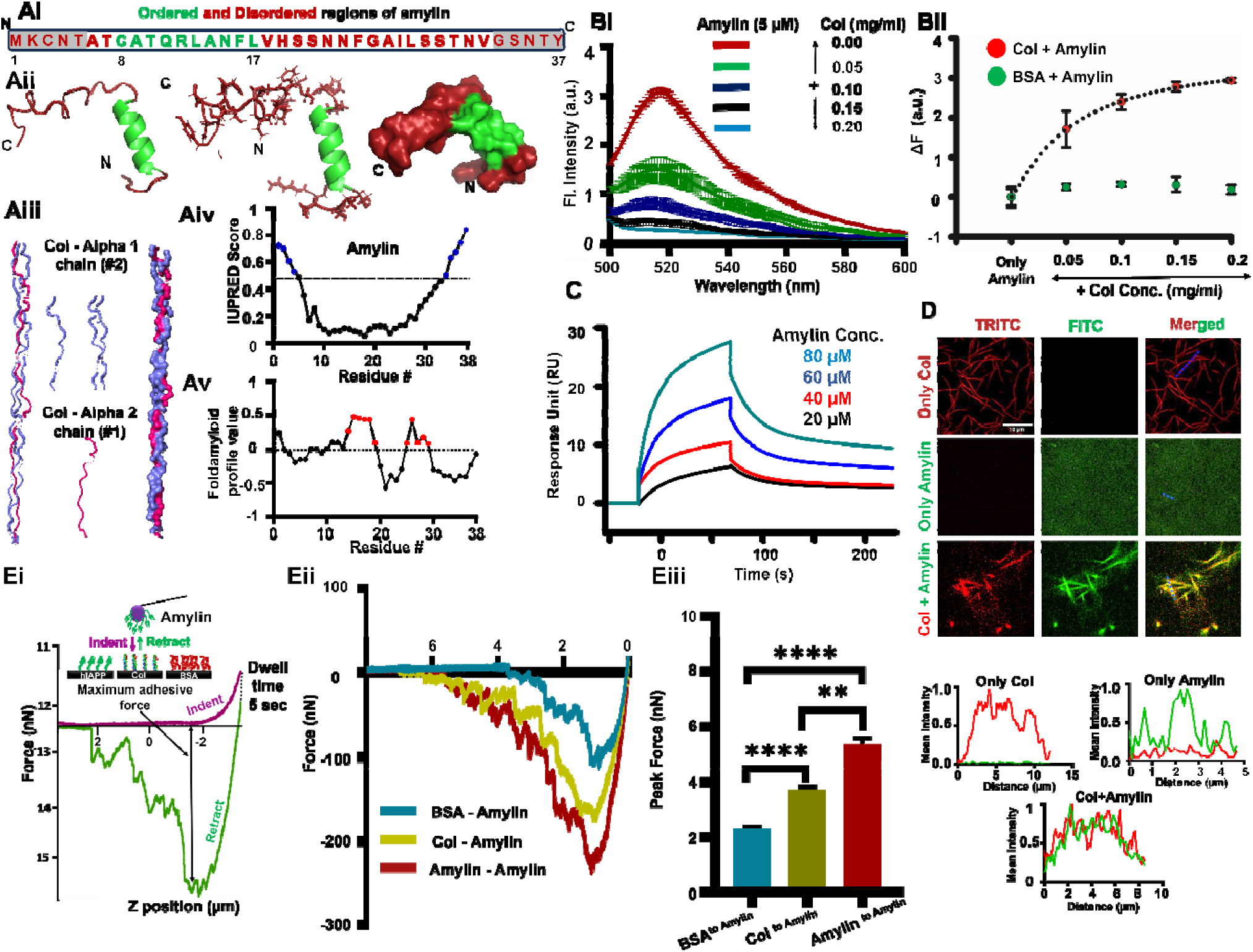
Amylin interacts with Collagen I (Col I). (Ai) Amino acid sequence of amylin (37 amino acids) showing ordered (8-17 aa, green) and disordered regions (1-7 & 18-37, red). (Aii) Three-dimensional (3D) structure of amylin showing its unstructured (red) and helix regions (green) (PDB ID: 5MGQ)^103^ (upper panel). (Aiii) 3D structure of Col I showing interwinding of 2 chains (blue) and 1 chain (pink) to form triple helical monomer (3HR2)^79^. (Aiv) IUPRED scores of amylin amino acids with red regions corresponding to intrinsically disordered segments. (Av) Aggregation-prone residues (red) of amylin mapped using Foldamyloid algorithm. (Bi) Representative fluorescence spectra of 5 µM FITC-amylin in the absence and presence of increasing concentrations of Col I. (Bii) Binding isotherm of 5 µM FITC-amylin in the absence and presence of increasing concentrations of Col I, with BSA serving as negative control ( samples per condition pooled from independent experiments). Error bars represent SEM. (C) SPR sensogram showing the kinetics of interaction of Col I with different concentrations of amylin. (D) Representative image showing colocalization of FITC-amylin (green) with Col I fibrils (red). (Ei) Quantification of strength of adhesion between amylin and Col I fibrils using Atomic Force Microscopy (AFM). Amylin/Col/BSA coated coverslips were indented with amylin-functionalized spherical probes. After indentation, tip was retracted after 5 sec dwell. (Eii) Representative retraction force curves obtained by indenting amylin/Col/BSA coated coverslips with amylin-functionalized probe. (Eiii) Quantification of maximum adhesive force or Peak Force (n 2: 30 force curves per condition pooled from N = 3 independent experiments) Statistical significance was assessed using One-Way ANOVA or nonparametric test. (** p < 0.01,**** p < 0.0001, ns: not significant).

To measure the strength of interaction between amylin and Col I, we performed adhesion experiments using AFM. Specifically, amylin functionalized spherical probes were brought in contact with BSA, Col I and amylin-coated glass coverslips to enable adhesion formation, and then retracted. The peaks observed during tip retraction correspond to breaking of these adhesion bonds (Fig. 2E). Since the shape of the retraction curves varied considerably between the conditions with the presence of multiple peaks, only the maximum unbinding force (or peak force) was compared. Compared to an average peak force of 2 nN for amylin-BSA interactions, the peak force of 3.5 nN observed for amylin-Col I interaction is indicative of a nearly two-fold stronger interaction, but weaker than that between amylin biding to itself (5 nN). Together our findings suggest that amylin is capable of binding to both Col I monomers and Col I fibrils with a moderate strength of interaction (Fig. 2F).

### ^15^N-^1^H HSQC and CPMAS NMR spectra reveal binding sites over amylin and Col I surfaces respectively during amylin-Col I interaction

In order to understand the mechanistic details by which collagen I interacts with amylin and initiates amyloid formation, we used solution NMR methods, which provide a comprehensive set of approaches able to characterize residue-specific features of strong protein−protein interactions on various time scales (52−55). A titration of Col I (0.2−0.8 mg/mL) with amylin monomer (100 µM) solution showed no significant chemical shift perturbations (except residue no 2, 10, 18, 21, 28, 37) in ^15^N-^1^H heteronuclear single quantum correlation (HSQC) spectra (Fig. 3A, Bi-ii, Ci & iii). However, a global residue-specific attenuation of the peak intensities was observed with increasing Col I concentrations (Fig. 3A-Bi, Supp. Fig. 2), suggesting strong amylin−Col I interactions (formation of a high molecular weight complex and subsequent decrease of NMR observable population of monomeric amylin). To minimize Col I aggregation during the NMR experiments and to capture the most specific interactions, we proceeded with low Col I concentrations (0.2−0.8 mg/mL) that displayed consistent residue-specific perturbations and kept samples at 10 °C. Under these conditions, the addition of 0.8 mg/mL Col I to amylin resulted in a reduction in resonance intensity of all peaks, consistent with transient formation of a high molecular weight complex (Fig. 3A-Bi). However, the greatest reduction in peak intensities occurred for residues 13 & 18-37 of the wild-type protein, which could be the anchor point of the amylin-Col I interaction (Fig. 3B).

**Figure 3:**
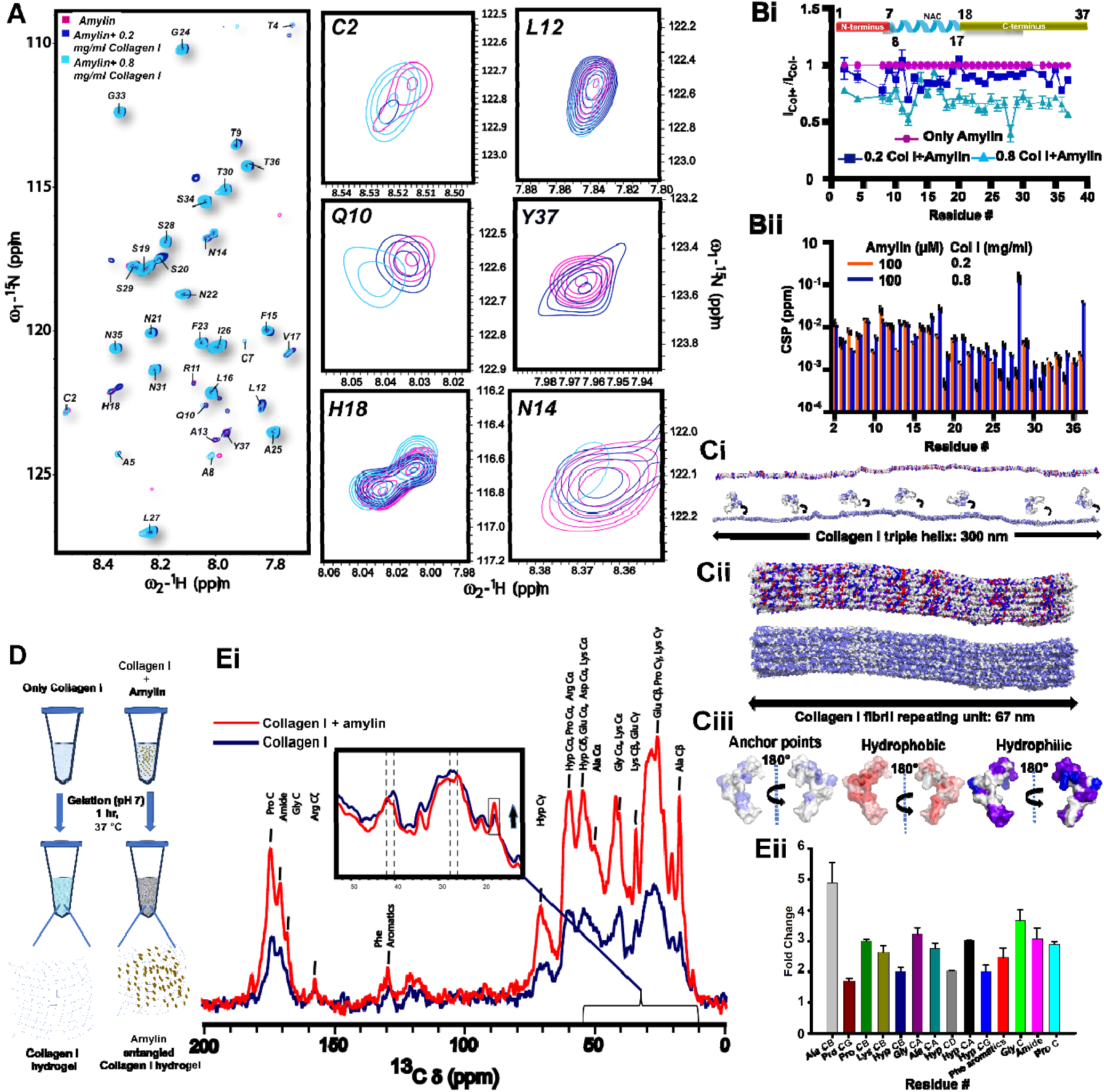
Characterization of residue-specific amylin−Col I binding using NMR. (A) ^1^H−^15^N-HSQC spectra of the amylin and Collagen I (0.2 or 0.8 mg/ml) incubated amylin shows global peak intensity loss of amylin upon collagen I addition. The inset shows a zoom-in on the 2D contours of residues that have higher peak shifting (Cys 2, Gln 10, His 18, leu 12, Tyr 37, Asn 14) upon addition of collagen I. (B) Amide backbone signal intensity ratios from ^1^H−^15^N-HSQC spectra of 100 μM amylin in the presence of 0.2 mg/mL (blue) & 0.8 mg/ml (indigo) collagen I compared with values in the absence of collagen I (pink). Dips in the signal intensity reflect regions maximally perturbed by the presence of collagen I. Noise levels are propagated as error bars in the plot. The secondary structural elements of amylin are indicated above the plot. (B) Cumulative secondary chemical shifts of amylin calculated from NMR backbone assignments. No significant chemical shift perturbation except residue no 18, 21, 28, 37. (C) (i) Surface model of the collagen I monomer (PDB: 3HR2)^79^ color-coded by amino acid type (top, hydrophilic; bottom, interacting residues based on ss-NMR data). A surface representation of amylin is shown for size comparison (PDB: 5MGQ)^103^. (ii) Surface models of the collagen I fibril repeating unit (built from PDB: 3HR2)^79^, color-coded by amino acid type (top, hydrophilic; bottom, interacting residues based on NMR data). The length of repeating unit is ∼67 nm, alongside mature fibrils can be ∼500 nm in diameter and microns long. (iii) Surface of amylin monomer (PDB: 5MGQ)^103^ color-coded by amino acid type (left, anchor points based on NMR data; middle, hydrophobic; right, hydrophilic). All models are color-coded as slate-interacting residues; red-acidic; blue-basic; purple-uncharged-polar; and orange, hydrophobic. (D) Cartoon shows gelation of collagen I and amylin incubated collagen I at pH 7 and 37 °C for solid state NMR experiments. (Ei-ii) Assigned resonances in natural-abundance 1D 1H−13C CPMAS spectra of Collagen I hydrogel (blue) and amylin incubated Collagen I hydrogel (red) shows peak intensity has been increased upon amylin addition. Peak intensity ratio has been plotted across different amino acids of Collagen I (right panel).

To monitor the interacting sites of Col I for amylin, we have performed Cross polarization magic Angle Spinning (CPMAS) experiment of Col I hydrogel vs amylin incubated Col I hybrid gel prepared by fibrillation/gelation of Col I in the presence/absence of amylin at pH 7-7.4 under 37 °C (Figure 3D). The intensity of the CPMAS spectra of Col I fibril increased significantly in addition of amylin suggests the orderedness of the Col I fibril increases after addition of amylin, and amylin shows a mixed pattern of interaction by associating with Col I through repeating hydrophobic (Hyp, Pro, Gly, Ala) and hydrophilic (Glu, Lys, Asp) residues as peaks of the same have been shifted upon presence of amylin in the hybrid gel (Figure 3Ci-ii, Ei-ii). Quantification of residue-specific fold change conveys a better clarity of Col I orderedness upon amylin binding where most of the residues show an average fold change of ≥ 2 (Figure 3Eii, Supp. Fig. 2). Collectively, the comparison of the molecular dimensions of interacting molecules—specifically, amylin (4 nm × 2 nm × 2 nm), Col I triple helix (300 nm × 1.5 nm), and mature Col I fibrils (microns in length and up to 500 nm in diameter)—demonstrates the potential for diverse binding modes. This diversity facilitates the independent binding of multiple β2m molecules to a single Col I molecule (Fig. 3Ci-ii). Notably, the surface of the Col I triple helix features a distribution of both hydrophilic and hydrophobic residues along its length (Figure 3Ci, Supp. Fig. 2). This surface chemistry pattern is retained on the Col I fibril surface (Fig. 3Cii, Supp. Fig. 2), further increasing the potential for various binding modes with complementary surfaces on amylin (Figure 3Ciii).

### MD simulations reveal strongest interaction between amylin fibrils and Col I fibrils

The stability of the simulated complexes was measured by root mean square deviation (RMSD) of the backbone atoms relative to initial structure and the compactness of the protein structure was studied by monitoring Radius of gyration (Rg) values (Figure 4). In the amylin monomer (AM)-amylin monomer complexed structure, the RMSD value ranges from 0.20 to 0.40 nm, and the Rg (radius of gyration) is between 1.25 to 1.35 nm. When AM is complexed with amylin fibril (AF), the RMSD decreases to 0.10 to 0.14 nm, while the Rg increases to 2.09 to 2.12 nm (Figure 4A-B). This indicates that the fibrillar forms are more stable compared to the monomer forms in terms of deviation. In the AM-AM complexed structure, large number of hydrophobic interactions were observed at the protein interface reflecting low RMSD values. Specifically, the distances between hydrophilic and hydrophobic residues fall within the range of 2.88 to 3.09 Å and 3.75 to 3.95 Å respectively. Conversely, in the AM-AF complexed structure, hydrophobic interactions between interface residues were coupled with C-H•••π interactions are observed (Table 2, Figure 4) where the distances range from 3.92 to 4.2 Å which provide stability to this complex. For AM complexed with Col I monomer (CM), the RMSD and Rg values vary from 0.20 to 1.0 nm and 2.12 to 2.20 nm, respectively, indicating a broader range of deviation compared to the AM-AF complex. When CM is complexed with AF, the RMSD ranges from 0.20 to 0.60 nm, and the Rg ranges from 2.20 to 2.28 nm. Notably, the binding of AM and AF to the CM/CF appears less stable in comparison to their interaction with the Col I fibril due to the limited surface area available for binding with a smaller number of interactions (Figure 4). In the Col I fibril (CF) complexed with AM, the simulated structures show an RMSD of 0.10 to 0.15 nm and an Rg ranging from 4.65 to 4.66 nm. When CF was simulated with AF, the RMSD and Rg values were observed to range from 0.15 to 0.30 nm and 4.77 to 4.81 nm respectively (Figure 4A). These observations suggest that the complexes involving amylin fibrils tend to exhibit lower deviation and higher stability compared to those involving amylin monomers. The AF molecules provide a considerable surface area for AM binding thereby increasing the stability of the complex. Furthermore, the AF complexed with CF exhibits the highest stability, attributed to the expansive surface area for AF binding on Col I. This configuration facilitates a large number of hydrophilic and hydrophobic interactions at the interfaces consequently enhancing their stability (Figure 4).

**Figure 4.**
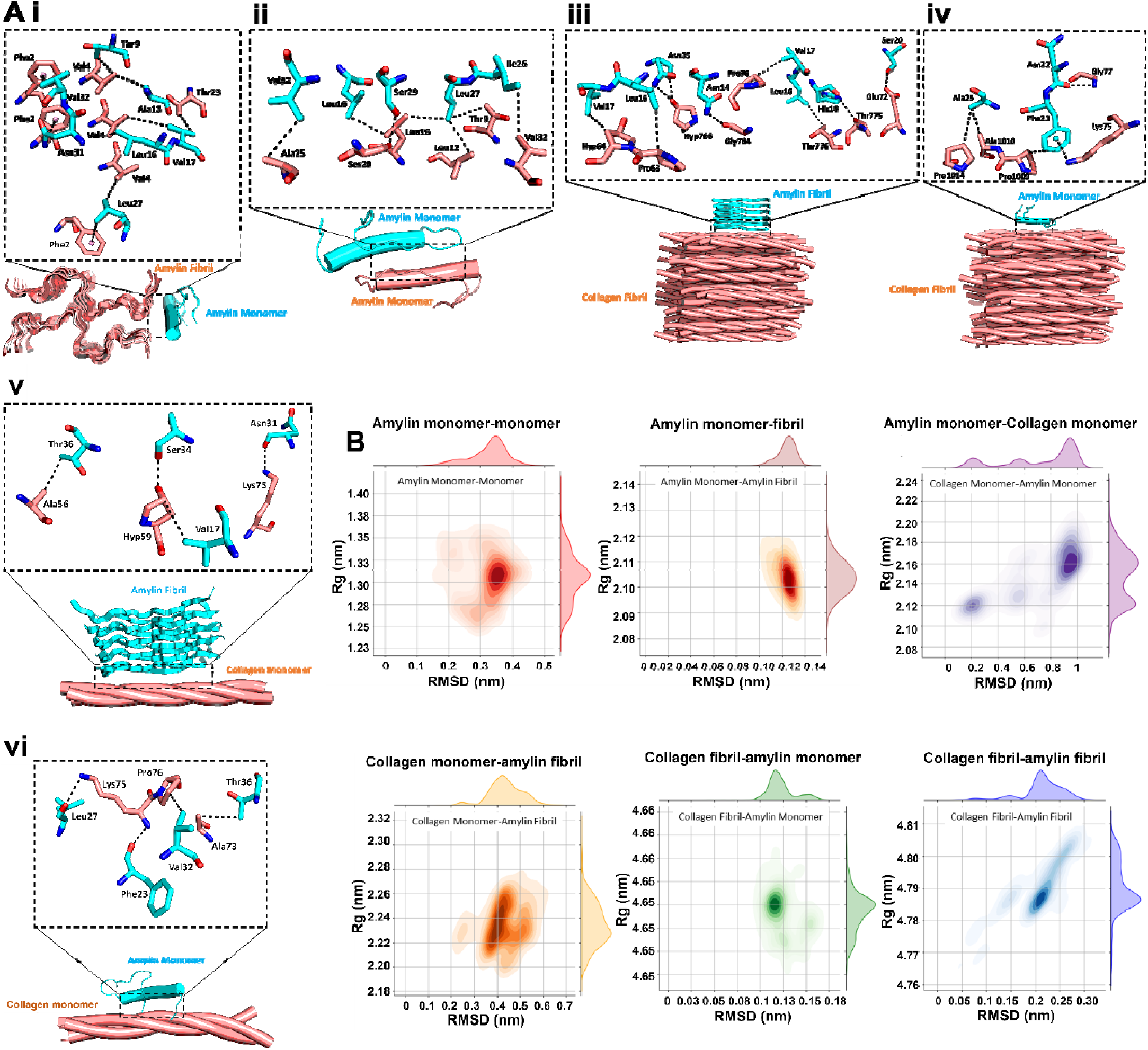
MD simulations depicting interactions between amylin monomers/fibrils with Col monomers/fibrils. (A) Interactions between different species (i.e., amylin monomers with amylin fibril (i), amylin monomers with monomers (ii), amylin fibril with collagen fibril (iii), amylin monomer with collagen fibril (iv), amylin fibril with collagen monomer (v) and amylin monomer with collagen monomer (vi)) obtained using MD simulations. Interactions between residues are shown by dotted lines. (B) Root mean square deviation (RMSD) vs radius of gyration (Rg) curves for the simulated complexes.

**Figure 5:**
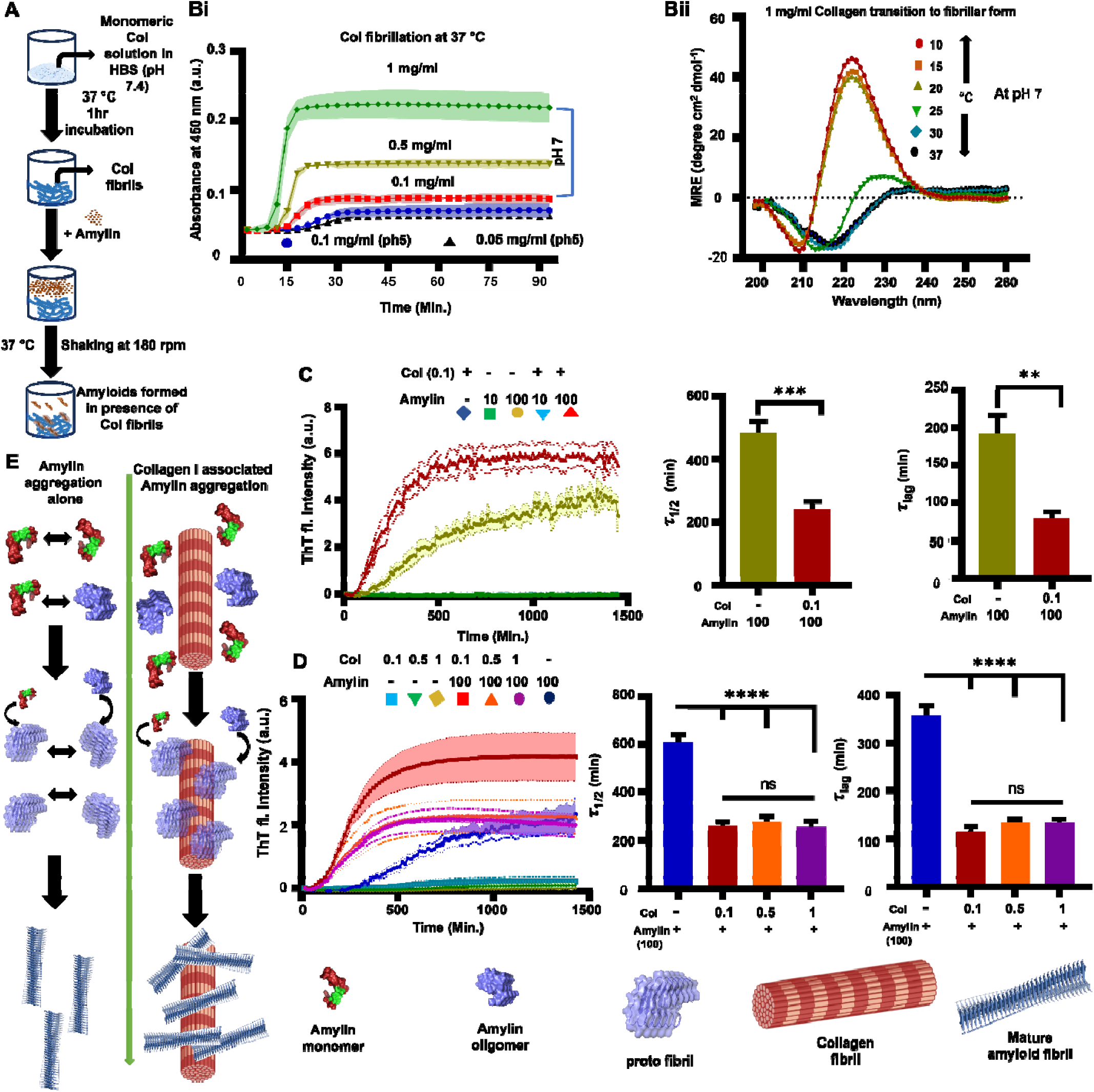
Influence of Col I fibrils on amylin aggregation. (A) Experimental setup for Col fibril preparation and subsequent amylin incubation for aggregation. (Bi) Concentration-dependent Col fibrillation at 37 °C and pH 7 & 5 (left panel) assessed using absorbance. (Bii) Self-assembly of Col tracked by measuring the mean residual ellipticity (MRE) as a function of wavelength at different temperatures. (Ci-iii) Aggregation of amylin at different concentrations tracked using ThT fluorescence in the presence and absence of Col fibrils. ThT curves which exhibited saturation were fitted to determine **_1/2_** and **_lag_** respectively ( samples per condition pooled from independent experiments). Error bars represent ± SEM. Statistical significance was assessed using unpaired t-test (**p < 0.01, ***p < 0.001). (Di-iii) Aggregation of 100 µM amylin at different concentrations of Col fibrils tracked using ThT fluorescence. ThT curves which exhibited saturation were fitted to determine T**_1/2_** and T**_lag_** respectively (n = 6 samples per condition pooled from N = 3 independent experiments). Error bars represent SEM. Statistical significance was assessed using One-Way ANOVA or nonparametric test. (****p < 0.0001, ns: not significant). (E) Schematic showing model of amylin aggregation in the presence and absence of Col fibrils.

**Table 2:** Hydrophobic, hydrophilic and π-interactions at the protein-protein interface of different combinations assessed using MD simulations.

| | Hydrophobic interaction | Hydrophilic interaction | π-interaction |
| --- | --- | --- | --- |
| Amylin Monomer-Amylin Monomer | Ala25<sub>CB</sub>•••Val32<sub>CG1</sub><br>Leu16<sub>CD1/CD2</sub>•••Leu16<sub>CD1</sub><br>Leu16<sub>CD2</sub>•••Leu27<sub>CD1</sub><br>Leu12<sub>CD1/CD2</sub>•••Leu27<sub>CD1</sub><br>Thr9<sub>CG2</sub>•••Leu27<sub>CD1</sub><br>Val32<sub>CG1</sub>•••Ile26<sub>CD1</sub> | Ser28<sub>OG</sub>•••Ser29<sub>OG</sub> | - |
| Amylin Fibril-Amylin Monomer | Val4<sub>CG1</sub>•••Thr9<sub>CG2</sub><br>Val4<sub>CG1</sub>•••Ala13<sub>CB</sub><br>Thr23<sub>CG2</sub>•••Val17<sub>CG1</sub><br>Val4<sub>CG1</sub>•••Val17<sub>CG2</sub><br>Val4<sub>CG2</sub>•••Leu16<sub>CD2</sub><br>Val4<sub>CG2</sub>•••Leu27<sub>CD1</sub> | - | Phe2($\pi$)•••Leu27<sub>CD2</sub><br>Phe2($\pi$)•••Asn31<sub>ND2</sub><br>Phe2($\pi$)•••Val32<sub>CG1/CG2</sub> |
| Amylin Monomer-Collagen Fibril | Ala25<sub>CB</sub>•••Ala1010<sub>CB</sub><br>Ala25<sub>CB</sub>•••Pro1014<sub>CG</sub><br>Phe23<sub>CE1</sub>•••Pro1009<sub>CB</sub> | Gly77<sub>NB</sub>•••Asn22<sub>OB</sub> | Phe23($\pi$) •••Lys75<sub>NZ</sub> |
| Amylin Fibril-Collagen Fibril | Val17<sub>CG1</sub>•••HYP<sub>CD</sub><br>Leu16<sub>CD1</sub>•••Pro63<sub>CA</sub><br>Val17<sub>CG1</sub>•••Pro76<sub>CG</sub><br>Leu16<sub>CD2</sub>•••Thr776<sub>CG2</sub> | Asn35<sub>ND1</sub>•••HYP<sub>OH</sub><br>Asn35<sub>NE2</sub>•••HYP<sub>OH</sub><br>Asn14<sub>NE2</sub>•••Gly764<sub>OB</sub><br>His18<sub>NE2</sub>•••Thr775<sub>OG1</sub><br>Ser20<sub>OG</sub>Glu72<sub>OE2</sub> | - |
| Amylin Monomer-Collagen Monomer | Val32<sub>CG1</sub>•••Pro76<sub>CG</sub><br>Thr36<sub>CG2</sub>•••Ala73<sub>CB</sub> | Leu27<sub>OB</sub>•••Lys75<sub>NZ</sub><br>Phe23<sub>OB</sub>•••Lys75<sub>OB</sub> | - |
| Amylin Fibril-Collagen Monomer | Thr36<sub>CG2</sub>•••Ala56<sub>CB</sub><br>Val17<sub>CG1</sub>•••HYP59<sub>CG</sub> | Ser34<sub>OG</sub>•••HYP59<sub>OG</sub><br>Asn31<sub>OB</sub>•••Lys75<sub>NZ</sub> | - |

#### Col I fibrils hasten amylin aggregation

Having established robust interaction between amylin fibrils and Col I fibres using MD studies, we next aimed to experimentally investigate the role of Col I monomers versus fibres on amylin aggregation kinetics (Fig. 4Ai, ii). For doing this, experimental conditions for obtaining Col I fibres were optimized by keeping the temperature fixed at 37 °C and varying pH and monomer concentration. Turbidity assay revealed concentration-dependent increase in fibrillation at pH 7, with negligible fibrillation at pH 5 (Fig. 4C, Supp. Fig. 3). To monitor Col I structural transitions, we performed temperature-sensitive circular dichroism (CD) analysis, which revealed structural changes in Col I at pH 7 with increasing temperature (Fig 4D). We observed a transition from the monomeric triple helix to the fibril form as the temperature reached 25°C, with further progression in fibrillation upon further temperature increase. Fibrillation appeared to saturate with time at temperatures above 25°C (Fig 4D). Transition to fibrillar state was not observed at pH 5 (Supp. Fig. 3).

To monitor the effect of Col I monomers and fibrils in regulating amylin aggregation, we incubated 10 & 100 µM amylin in the presence and absence of 0.1 mg/ml Col I fibrils formed at pH 7, 37°C. Strikingly, Col I fibrils hastened aggregation of 100 µM amylin (monitored by ThT intensity) as observed from the significant reduction in r_l/2_ & r_lag_. However, Col I fibrils did not hasten aggregation of 10 µM amylin (Fig 4Ei). To check the effect of fibril density on amylin aggregation, 100 µM amylin was incubated with varying concentrations of Col I fibrils (0.1, 0.5 & 1 mg/ml). We saw a similar pattern of hastened amylin aggregation across all Col I concentrations with no significant changes with respect to Col I fibrillar density (Fig 4Eii). Collectively, our results demonstrate the role of fibrillar Col I in enhancing amylin aggregation.

### Amyloid entangled Col I matrices (AEMs) are toxic to islet **β** cells

Robust interaction between amylin and Col I raises the possibility that in vivo, to study the implications of such amylin-enriched ECMs on f3 cell viability and functionality, INS1E-β cells were cultured on amylin entangled Col I matrices (hereafter referred to as AEMs) (Fig. 6Ai). In AEMs generated using 100 µM amylin (i.e., AEM100), amylin formed amyloid aggregates as evidenced from the co-localization of amylin (red) with amyloid (green) and Col I fibrils (grey) (Fig 6Aii). To evaluate the impact of such matrices, INS1E-β cells were seeded on AEM10 and AEM100 matrices, with Col I gels and coverslips incubated with amylin (10 and 100 µM) (i.e., Amylin10, Amylin100) serving as controls. Remarkably, compared to controls, both islet size and the number of cells per islet were significantly diminished in cells cultured on AEM100 substrates (Fig 6Bi-ii). This was associated with a reduction in cell surface area, nucleus size, and integrin β1 expression (Supp. Fig. 4).

**Figure 6.**
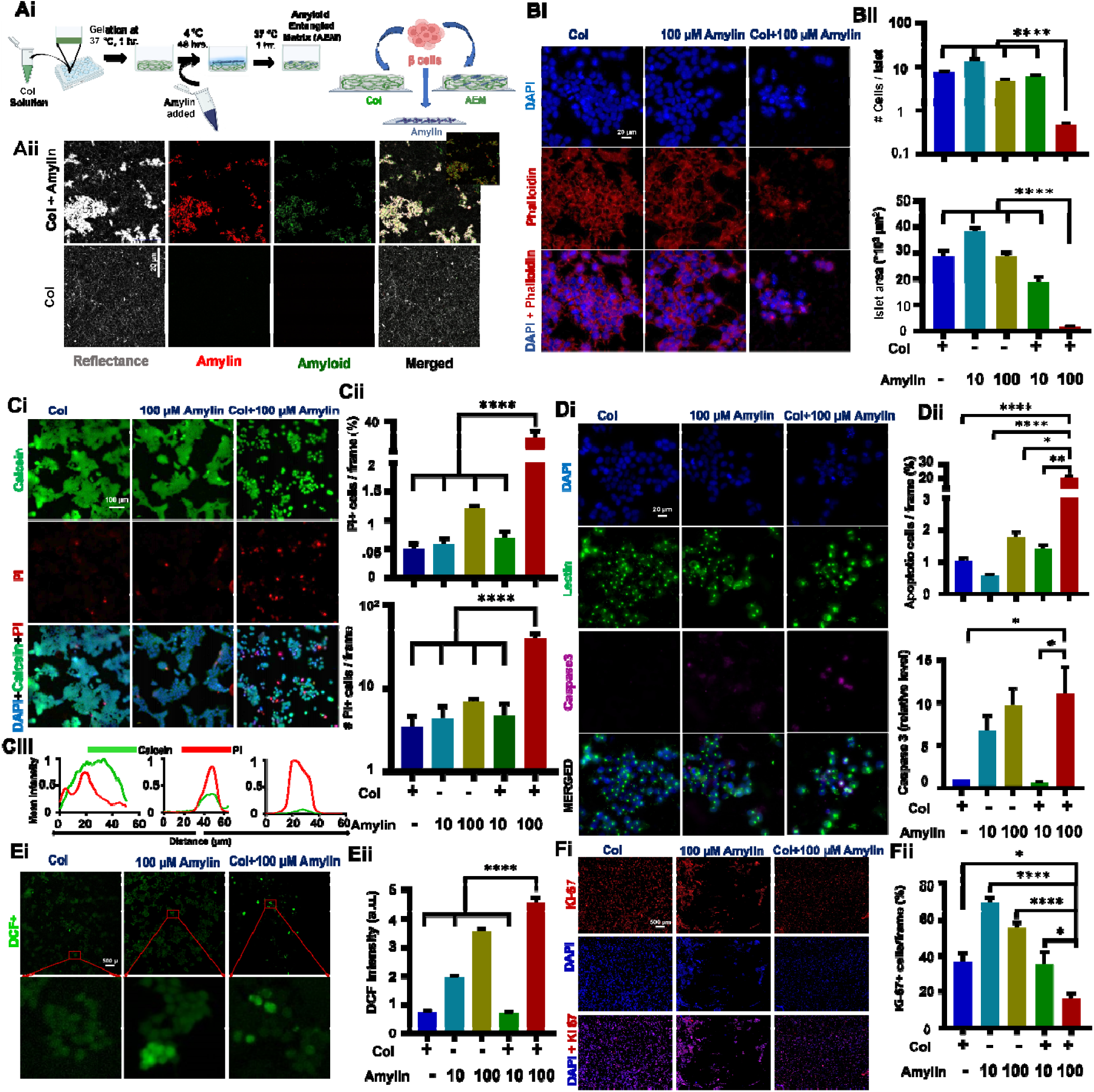
Amyloid entangled matrices (AEMs) containing high amylin concentration are toxic to INS1E. β **cells.** (Ai) Experimental setup for preparing AEMs and culturing of β cells on AEMs. (Aii) Representative images of Col hydrogels and AEMs obtained by combining reflectance with immunofluorescence microscopy. Samples were co-stained with amylin antibody (red) and amyloid-specific OC antibody. (Bi-ii) Representative phalloidin/DAPI-stained images of INS1E β cell after 48 hr culture on Col gels, amylin coated coverslip & AEM100 (AEM obtained with 100 M amylin), and quantification of islet morphology (≈70 - 90 cells per condition, N = 3). Scale bar 20 µm. (Ci-iii) Live-dead staining of INS1E β cells using calcein/PI and quantification of cell viability and mean intensity (≈15 frames per condition, *N* = 3). (Di) Representative lectin (green), caspase 3 (Pink) and DAPI (blue) stained images of INS1E β cell across different conditions, and quantification of fraction of Caspase 3+ cell/frame (≈15 frames per condition, *N* = 3) (right panel). (Dii) Caspase 3 gene expression in INS1E β cells across different conditions normalized to that on Col gels (*N* = 3 per condition). (Ei, ii) Representative image showing ROS expression in INS1E β cells using DCFDA assay and quantification of mean DCF intensity across different conditions (≈80-100 cells per condition, *N* = 3). (Fi, ii) Representative KI67 (red) and DAPI (blue) stained images of INS1E β cell across different conditions and quantification of fraction of KI67+ cell per frame (≈6 frames per condition, *N* = 3). Data are presented as mean ± SEM. Statistical significance was assessed using One-Way ANOVA or nonparametric test (* p < 0.05,** p < 0.01,*** p < 0.001,**** p < 0.0001, ns: not significant).

To elucidate the reason behind the reduction in islet size on AEM100, live cell staining with PI and Calcein AM was performed. Interestingly, compact islets (green-Calcein AM) with a limited number of PI+ cells were observed on Col I, Amylin 10, Amylin100 and AEM10 substrates; in contrast, islets were scattered on AEM100 with a significant increase in the proportion of PI+ cells (Fig 6Ci-ii). Moreover, most of the PI+ cells on AEM100 exhibited reduced Calcein AM intensity, indicative of membrane rupture as a possible mechanism of cell death on AEM100 matrices (Fig 6Ciii). Increase in caspase 3 intensity along with increase in transcript caspase 3 levels (Fig. 6Di-iii), as well as high levels of necroptotic markers RIPK3 and MLKL (Supp. Fig. 4) suggests that AEMs induce cell death via apoptosis and necroptosis. Since amylin oligomers and fibrils induce high ROS generation through induction of ER & mitochondrial stress in different β cell models ^51,68,69^, intracellular ROS levels were assessed using DCFDA assay, and revealed highest intensity in INS1E cells cultured on AEM100 matrices (Fig 6Ei, ii). Finally, assessment of cell proliferation using Ki67 staining revealed lowest proliferation levels in cells cultured on AEM100 matrices (Fig 6Fi-ii). Collectively, these results indicate AEM with high levels of entangled amylin induce f3 cell death and inhibit cell proliferation.

### AEM100 matrices perturb **β** cell functionality

To gain insight into the implications of amyloid-deposited Col I fibrils on β cell functionality, we investigated the glucose uptake ability of INS1E cells on these matrices. Therefore, we incubated the cells with 100 µM 2-NBDG for a time being and observed a decrease in 2-NBDG intensity, indicating reduced glucose uptake ability in INS1E cells present on Col I+100 Amylin compared to others (7Bi- ii). Subsequently, we sought to evaluate insulin expression of INS1E β cells using anti-Insulin antibodies under various conditions. Surprisingly, we observed higher expression of insulin (measured by mean insulin intensity in 70-90 cells per condition) in β cells seeded on AEM100 (Fig 7Ai-ii). As a supporting study of high intracellular insulin levels, we assessed the levels of insulin regulatory markers and found upregulation of MAFA, NKX6.1 and PAX6, indicative of increased transcription of the insulin gene, as they bind to insulin gene promoter to facilitate insulin production (7Aiii). To monitor glucose sensitivity and subsequent release of insulin, we performed glucose-stimulated insulin secretion, in which we initially incubated the cells at 2.8 mM glucose, and after a while, we increased the concentration of glucose to 16.7 mM. We found that insulin release has not been significantly increased upon (fold change 1.8±0.5) 16.7 mM glucose spike in INS1E present on AEM100 compared to only Col & AEM10, which indicates an impaired insulin release despite intracellular high insulin levels.

**Figure 7:**
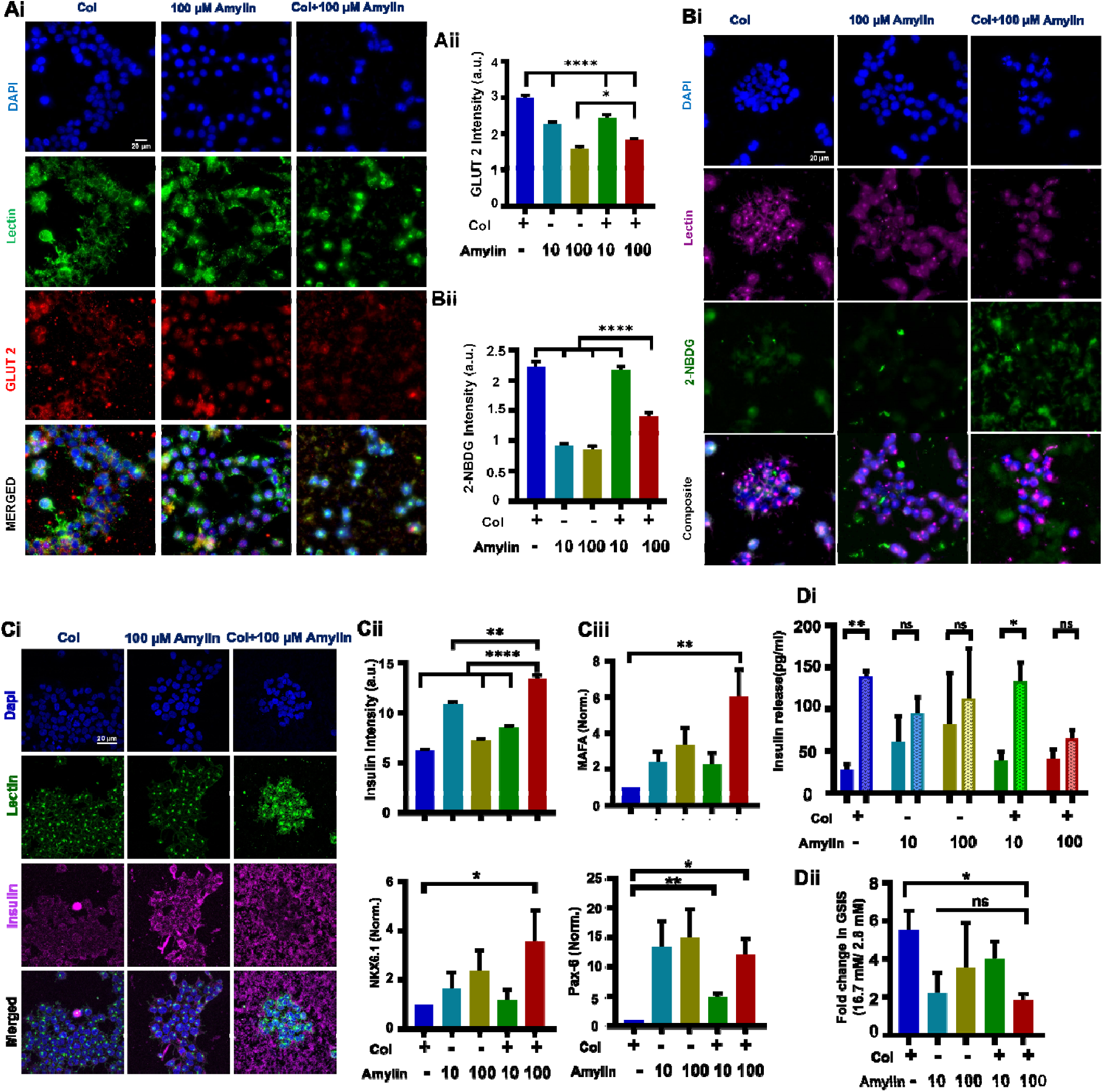
Amyloid entangled matrices (AEMs) containing high amylin concentration cause loss of. β **cell functionality.** (Ai, ii) Representative GLUT2 (red), lectin (green) and DAPI (blue) stained images of INS1E β cell across different conditions, and quantification of mean GLUT2 intensity ( cells per condition, ). (Bi, ii) Representative fluorescence images depicting 2-NBDG (green) uptake and its quantification (( cells per condition, ) in INS1E β cells across different conditions. Cells were co-stained with lectin (green) and DAPI (blue). (Ci, ii) Representative insulin (pink), lectin (green) and DAPI (blue) stained images of INS1E β cell across different conditions, and quantification of mean insulin intensity (≈ 80 - 100 cells per condition, N = 3). (Ciii) Relative mRNA expression of MAFA, NKX6.1 and PAX6 genes in β cells across different conditions (N = 3). (Di, ii) Glucose stimulated insulin secretion in INS1E cells cultured across different conditions, and quantification of fold-change (N = 3). For all sub-figures, data is presented as mean ± SEM. Statistical significance was assessed using Two-Way ANOVA &/or nonparametric test (* p < 0.05,** p < 0.01,*** p < 0.001,**** p < 0.0001, ns: not significant).

## Discussion

Clinically, Type 2 Diabetes (T2D) progression is linked to β cell dysfunction and loss, with patients unable to produce sufficient endogenous insulin to counteract insulin resistance and control blood glucose levels ^70–72^. Although the primary cause of β cell failure in T2D remains unknown, the accumulation of aggregated forms of the peptide hormone amylin in pancreatic islets of T2D patients is a likely contributor to decreased β cell function, mass, and early disease progression ^11,34,73,74^. Previous experiments have focused on the kinetics of amylin mediated amyloid formation in the presence of various molecules or its impact on islet cell health ^19,43,75^. However, the contribution of the islet microenvironment on amylin aggregation, and its implications vis-à-vis survival and functioning of f3 cells have not been adequately addressed. In this study, for the first time, we document a strong interaction between amylin and collagen I—one of the principal constituents of the islet microenvironment, with fibrillar collagen I mediating faster aggregation of amylin and amylin aggregates driving β cell death and loss of function (Fig. 8).

**Figure 8:**
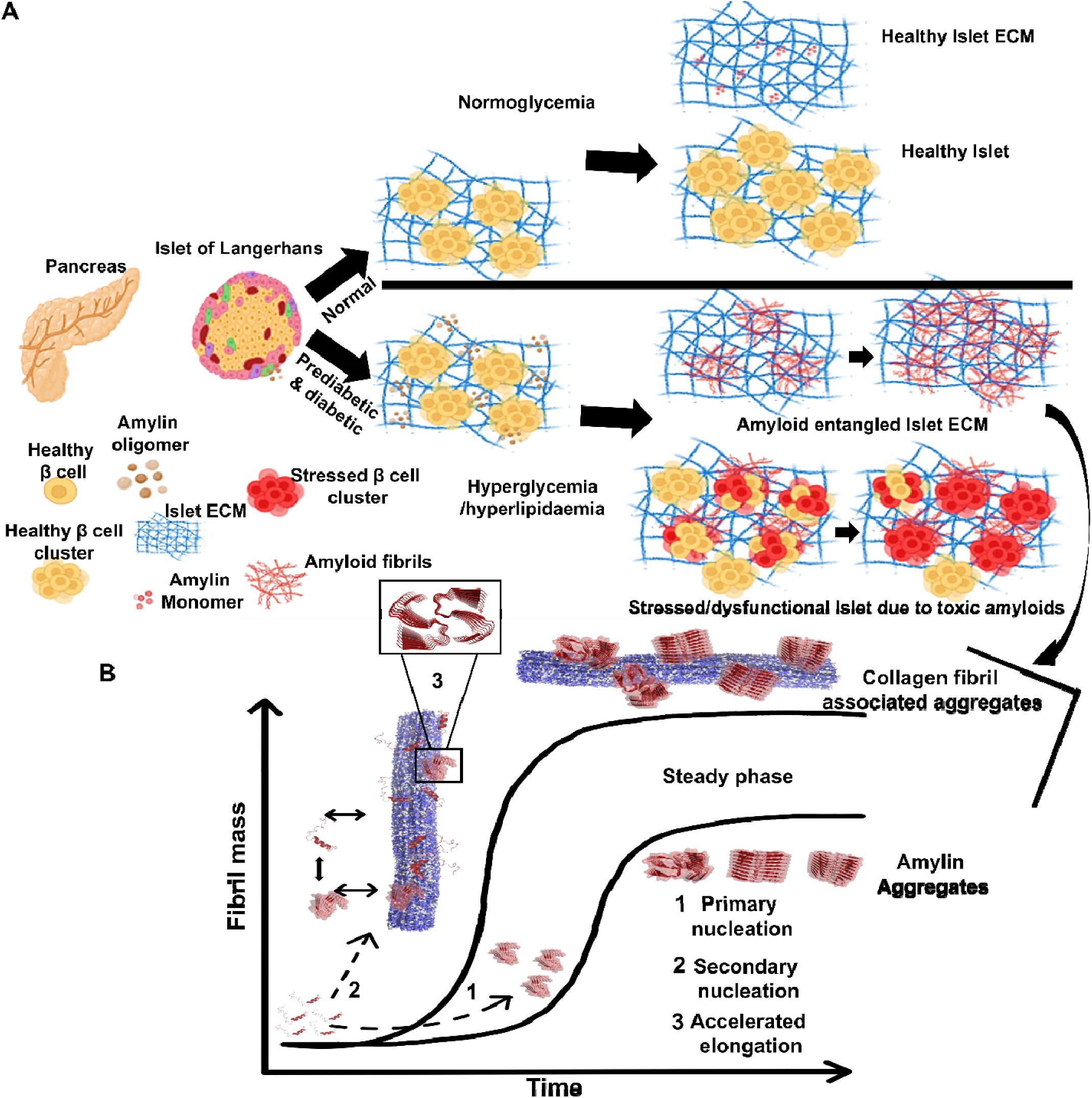
(A) Proposed model of amylin amyloid deposition in Islet tissues during diabetic progression. (Prepared from free version of biorender application) (B) Proposed model of amylin aggregation through surface templating by Col I.

Our fluorescence, SPR and AFM experiments establish a strong interaction between amylin and both monomer and fibrillar forms of collagen. A comparison of the molecular dimensions of the interacting molecules—amylin (∼ 4.6 nm **×** 1.6 nm **×** 1.6 nm), collagen I triple helix (300 nm **×** 1.5 nm **×** 1.5 nm), and mature collagen I fibrils (microns in length and up to 500 nm in diameter)—highlights the potential for a multitude of binding sites. This variability may thus enable multiple amylin molecules to independently bind to the same collagen molecule. Notably, the surface of the collagen I triple helix is interspersed with numerous hydrophilic and hydrophobic residues along its length. When collagen is in monomeric triple helical form, a greater fraction of surface exposed sites are hydrophobic. The hydrophobic nature of both amylin monomers and oligomers is thus likely to promote amylin interactions with collagen monomers ^76–80^. In line with this, the repeating pattern of hydrophilic and hydrophobic residues maintained on the collagen I fibril surface further enhances interaction with amylin ^36,79^.

GAGs such as HSPG and heparin have both been reported to accelerate amylin aggregation, and have been implicated in f3 cell toxicity ^19,27^. Both these molecules possess negative charge bearing surfaces which can interact with positively charged amylin (because of His) at physiological pH. Here, through NMR and MD simulation studies we show hydrophobic, hydrophilic and cation-π interactions collectively regulate amylin-collagen I interactions. Our NMR data suggests that amylin has few anchor points located towards the C terminus region, which facilitates its binding with collagen I. Apart from these, there are other binding sites distributed throughout its length that also help its complex formation with collagen I ^29,36,64,78,79^. Jean Baum and Sheena Radford’s groups have reported that as a lengthy protein, collagen I has multiple different binding sites for small globular protein such as β2 microglobulin with hydrophobic and hydrophilic interactions playing a major role in collagen I- β2 microglobulin complex formation and amyloid formation of β2 microglobulin ^29,36^. Likewise, our solid-state NMR data informs that collagen I has multiple binding sites (hydrophobic as well as hydrophilic in nature) for amylin, which are distributed throughout its length. Higher peak intensity of amylin incubated collagen I hydrogels compared to collagen I gels is indicative of lower flexibility of collagen I fibrils in the presence of amylin. This may occur due to binding of multiple amylin molecules on collagen I fibril surface making it bulky and hydrophobic. In vivo, these interactions may perturb islet ECM turnover by hindering the functions of collagenase and/or MMPs ^81^, inhibit cell-matrix communication by masking or competing for integrin binding sites ^82–85^, and also induce fibrosis by repelling water ^35,74,86^ leading to islet toxicity and decreased islet functionality ^87–91^.

Our MD simulations show that the complex formed between amylin monomers and amylin fibrils is more stable and expanded compared to the amylin monomer-monomer complex. On the other hand, either amylin monomer or amylin fibril complexed with collagen fibril shows a profile of reasonable stability similar to amylin monomer-fibril complexes with an expanded form of the structures because of more surface area for interactions over collagen fibril. Complexes that include either collagen and amylin monomers, or, collagen monomer and amylin fibrils, are relatively unstable due to a smaller number of interactions on the collagen monomer surface. Consistent with our MD simulations, our aggregation kinetics experiments show that the addition of collagen fibrils hastens amylin aggregation. Based on these findings, we speculate that the presence of collagen fibril is a potential source of stabilizing amylin monomer and/or fibrils over its surface, which are abundant in the diabetic islet. The fibrillar surface of collagen might function as a potential secondary nucleation site for amylin aggregation and lead to collagen-associated amyloid fibrils impacting islet endocrine cell health. Based on the RMSD values of different complexes, our findings suggest that expansion of amyloid deposits preferably occurs through addition of amylin monomers to existing amylin aggregates associated with collagen in comparison to addition over existing monomeric amylin bound to collagen fibrils. Since amylin is secreted under normoglycemic conditions also, we posit that amylin aggregation on collagen fibrils occurs only under hyperglycemic conditions when a threshold amylin concentration is reached. Given the role of fibrillar collagens in hastening amylin aggregation through surface templating, peptide or small molecule inhibitors that are able to block amyloid formation by preventing amylin-amylin interaction, may not be sufficient for clearance of ECM deposited amyloids as those drugs might not inhibit amylin-ECM interaction ^92–94^.

*In vivo* studies have shown that during diabetes, amyloid deposition eradicates insulin-positive β-cells from the islets, causing the islets to shrink ^87,89^. In vitro studies also report that amyloidogenic forms of amylin cause β-cell toxicity through ROS generation, creating ER and mitochondrial stress, plasma membrane rupture, and apoptosis ^7,51,63,68,90,95^. In line with this, presence of small, scattered cell clusters in AEM100 matrices and increased cell death may be attributed to increased amylin entanglement and amyloid deposition in in these matrices ^7,72,90^. In AEM100 matrices, collagen acts both as a driver of amyloid formation and as a substrate for their retention, paralleling the in vivo scenario where ECM remains associated with amyloids during diabetes. Consistent with previous reports from in vitro and humanized mouse models, we observed high intracellular ROS, low proliferation ability, and both necroptosis and apoptosis contributing to β-cell death on AEM100 substrates ^96^. To combat hyperglycemia, islets try to compensate by increasing β-cell number or insulin synthesis, but dysfunction and death of β-cells due to islet amyloid affect these compensatory mechanisms. In spite of higher insulin expression in β-cells observed on AEM100 substrates and in islet sections from diabetic mice, glucose-stimulated insulin release was impaired. Our findings suggest that the differential expression of insulin and defective insulin release in β-cells result from a multifactorial process involving ER stress, high intracellular ROS, and membrane rupture mediated by amyloids deposited in the matrix ^7,63,91^.

In conclusion, we have demonstrated for the first time that binding to collagen I may concentrate amylin molecules, thereby enhancing the likelihood of self-association or increasing the potential for secondary nucleation on the collagen I surface, both of which could promote fibril formation. These mechanisms, individually or collectively, might catalyse amylin aggregation in vitro, providing a molecular rationale for the deposition of amylin in collagen-rich islet tissues of diabetic patients. More broadly, these findings exemplify the critical role of the physiological environment in amyloid formation, explaining the specific deposition of amyloid in different tissues ^29,36,97^, and, in some instances, the deposition of different protein variants in different tissues and their pathogenicity ^97–102^. The methodologies employed here to study the strong interactions within the amylin-collagen I complex can be extended to future investigations. These studies can provide atomic-level insights into how other physiologically relevant cofactors promote amyloid formation in IDPs implicated in various amyloid diseases and how that such association impacts tissue health. Given the prominent role of collagen associated amylin aggregates in inducing β cell toxicity, targeting amylin-collagen and amylin-amylin interactions in combination may represent a prospective therapeutic approach for diabetes management.

## Supporting information

https://docs.google.com/document/d/1x43ZMevEW-QNdNbp13Z8aEPzmbHqTo0E/edit?usp=sharing&ouid=111200009834097026871&rtpof=true&sd=true

## Acknowledgements

We acknowledge IIT Bombay for providing Bio-AFM, NMR, CD, SPR and Confocal Microscopy facilities. We acknowledge Dr. Shilpi Sharma (Savitribai Phule Pune University) for providing INS1E β cell and its culture protocols. We acknowledge Zofishan Iqra Anjum, Md Wasim akram ddoza Hazari & for additional supports in data organisation.

## Author Contributions

Conceptualization: Md.A.H., A.K., S.Sen; Methodology: Md.A.H., F.S., K.P.M., S.Dg., S.P., S.S.; Formal analysis: Md.A.H., G.K., A.K.J., K.P.M., S.Dg., F.S., S.P., S.Dt., S.Sen.; Investigation: Md.A.H., A.K., S.Sen.; Data curation: Md.A.H.; Writing - original draft: Md.A.H., A.K., S.Sen; Writing - review & editing: Md.A.H., SK.D., S.P.P., H.Z., P.B., A.K., S.Sen; Supervision: A.K., S.Sen; Project administration: A.K., S.Sen; Funding acquisition: A.K., S.Sen

## Competing interests

The authors declare no competing or financial interests.

## Funding

S.Sen and A.K. acknowledge financial support from the Wadhwani Research Centre for Bioengineering, IIT Bombay (Grant # RD/0122-DONWR04-001, DO/2018-WRCB002-022). Md.A.H. was supported by the fellowship from the CSIR, Ministry of Human Resource Development India (Grant # CSIR 09/087(0950)/2019-EMR-I).

## Notes

### Competing Interest Statement

The authors have declared no competing interest.

