## Supplementary material for "Faster amylin aggregation on fibrillar collagen hastens diabetic progression through β cell death and loss of function": https://docs.google.com/document/d/1x43ZMevEW-QNdNbp13Z8aEPzmbHqTo0E/edit?usp=sharing&ouid=111200009834097026871&rtpof=true&sd=true

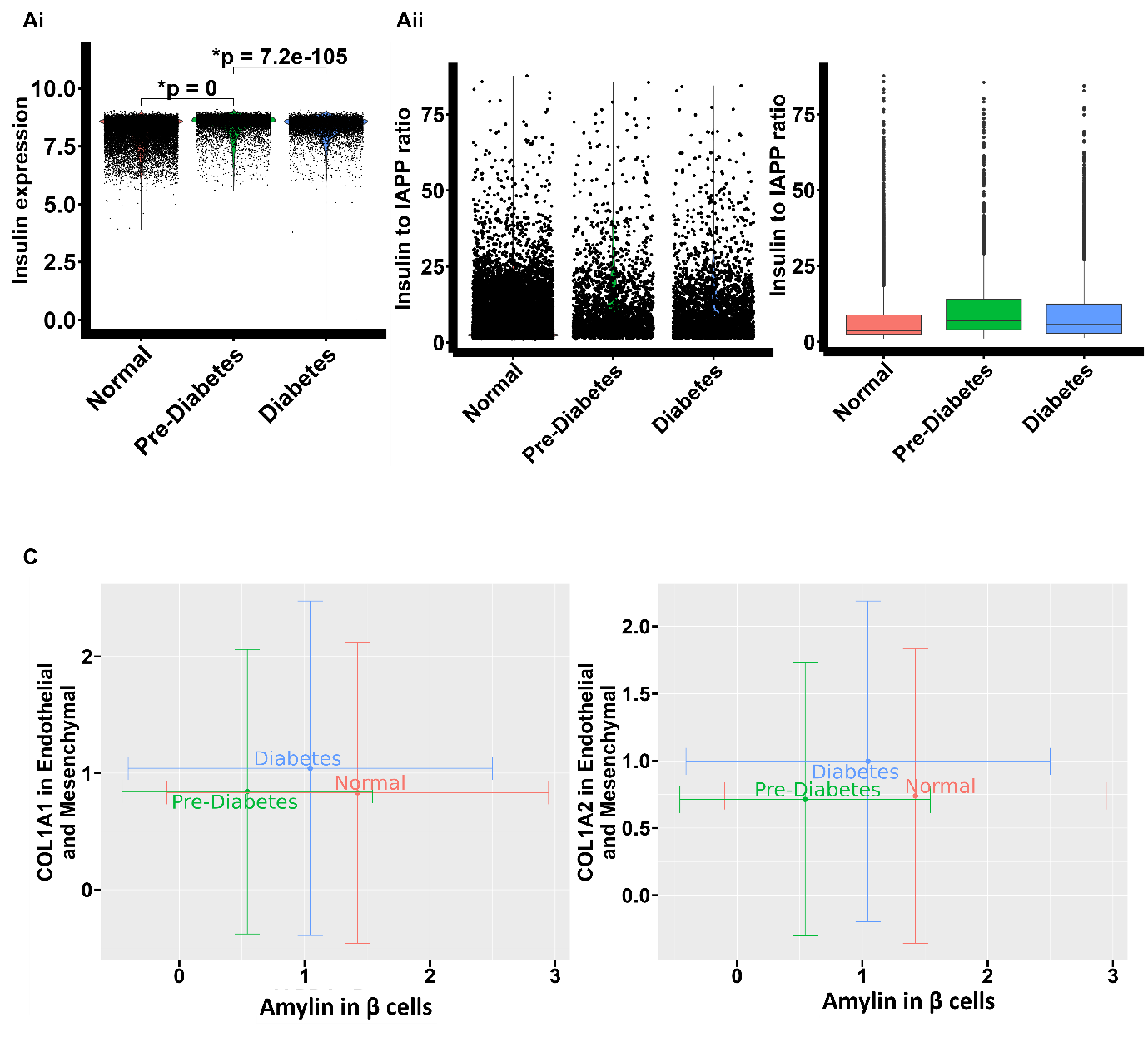

**Supp. Figure 1.** **Correlation of alterations in Collagen and amylin expression and their association during diabetes progression with islet organization and functionality.** (Ai-ii) The expression levels of insulin in β cells & ratio of insulin to amylin expressions in β cells is shown as volcano & box plot across normal vs prediabetic vs diabetic conditions. (Bi-ii) Correlation plot of amylin and Col I (Col IA1 &Col IA2) expression across normal vs prediabetic vs diabetic conditions.

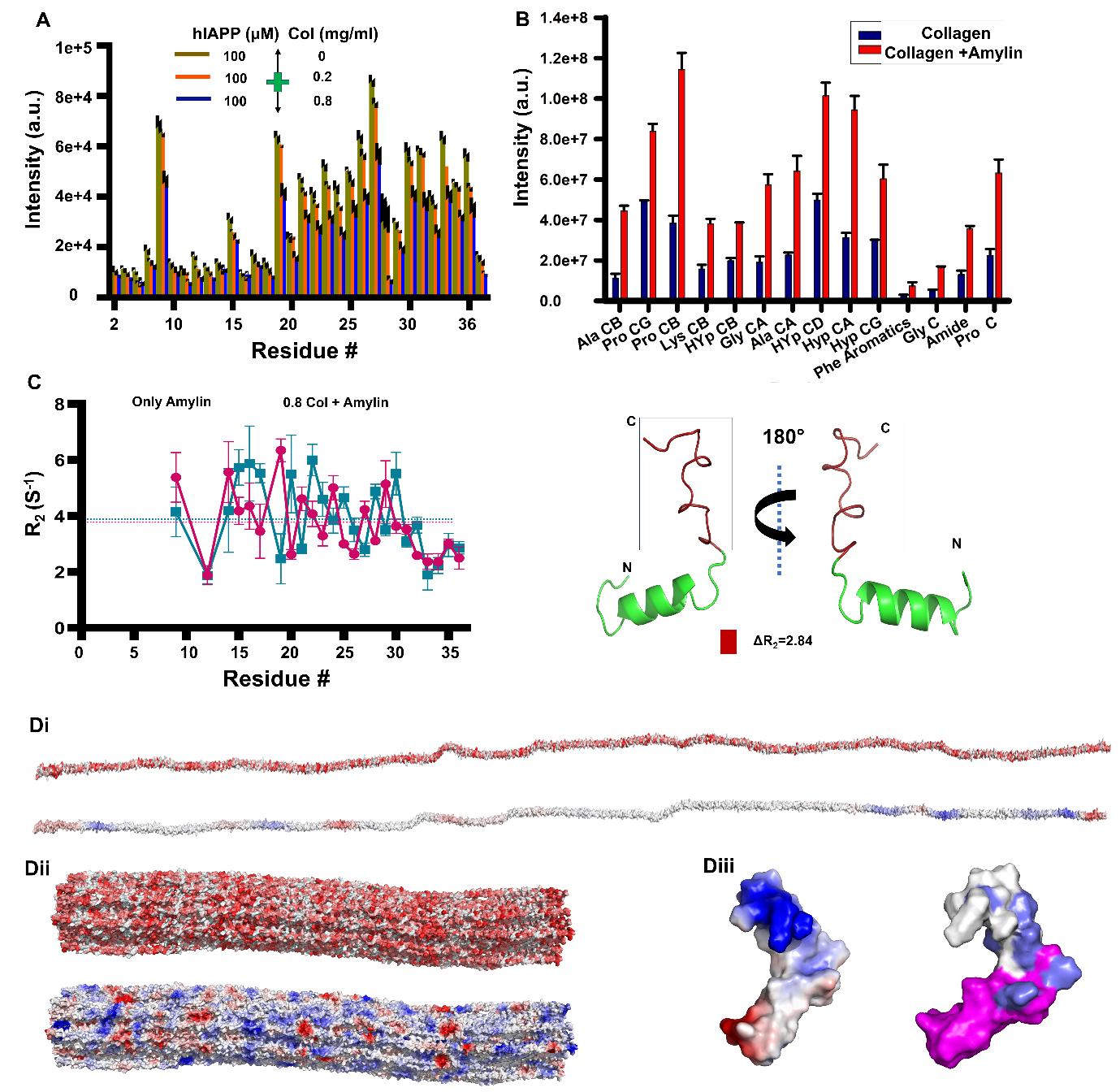

**Supp. Figure 2**. **Characterization of residue-specific amylin−Col I binding using NMR.** (A) Residue specific peak intensity derived from 1H-15N HSQC spectra of 100 µM amylin alone & in presence of 0.2 & 0.8 mg/ml Col I shows global decrease of peak intensity upon Col I incubation. (B) Residue specific peak intensity derived from Cross polarization magic Angle Spinning (CPMAS) experiment of Col I hydrogel and Col I + Amylin hybrid gel, shows rise in intensity upon amylin incubation in Col I hydrogel. (C) 15N-R2 measurements of 100 μM amylin in the presence (indigo) or absence (red) of 0.8 mg/mL Col I. The errors are propagated from the fitting errors. The dashed lines indicate the mean 15N-R2 values of amylin in the presence or absence of 0.8 mg/ml Col I. (Di-iii) Surface models of the collagen I triple helix (i) & fibril repeating unit (ii) (built from PDB: 3HR2) (Orgel et al., 2006) color-coded by amino acid type (top, hydrophobic; bottom, electrostatic). The repeating unit is ∼67 nm in length, however mature fibrils can be microns long and ∼500 nm in diameter. Distinct bands of electrostatic residues are observed within the repeating unit. (iii) Representative surface model of Amylin (built from 5MGQ) (Rodriguez Camargo et al., 2017) showing electrostatic surface profile and interacting residues (Slate: high chemical shift points, Pink: significant decrease in peak intensity).

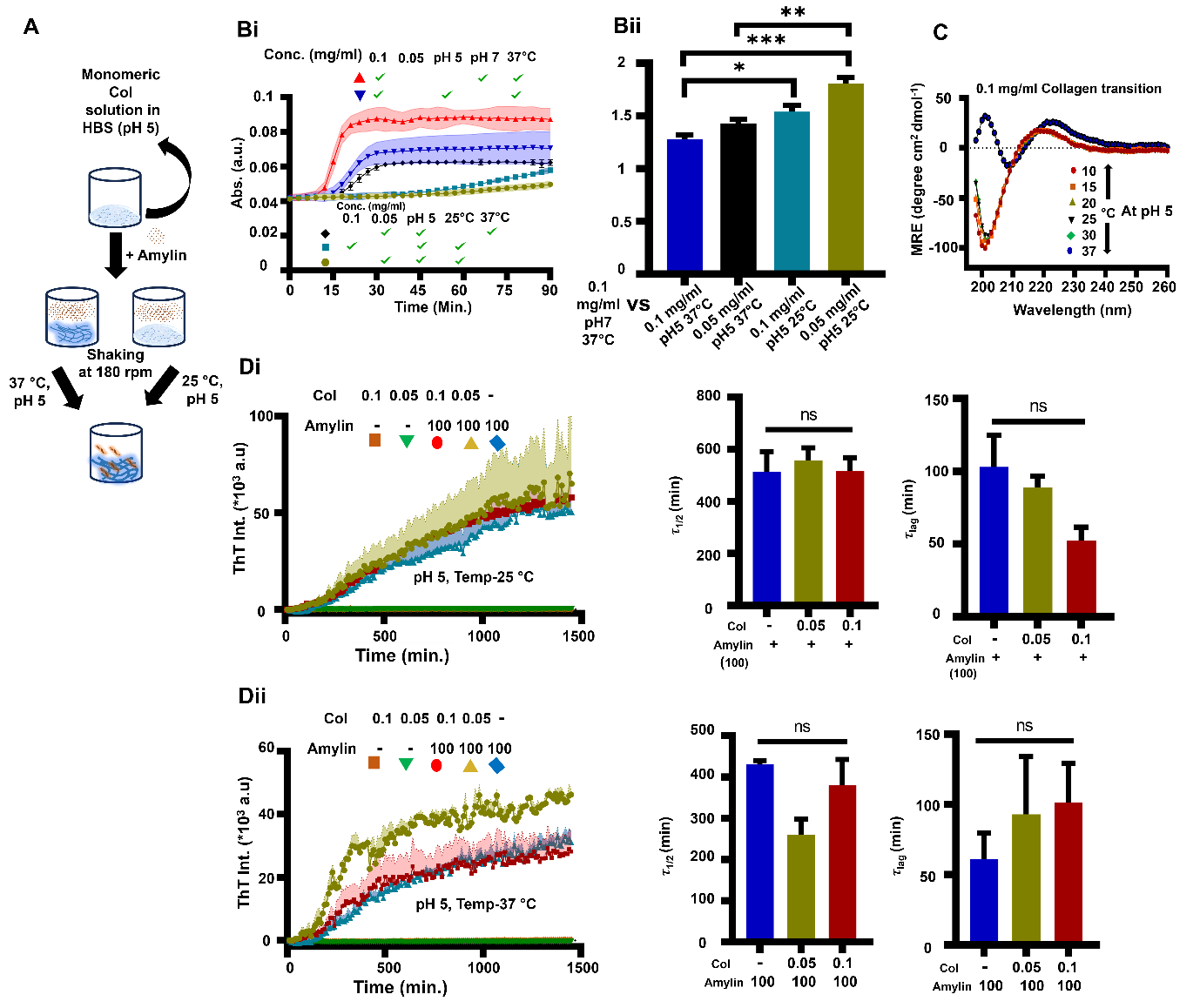

**Supp. Figure 3. Influence of Col I monomer on amylin aggregation.** (A) Representative experimental set up for amylin and Col I monomer aggregation. (Bi-ii) Representative figure shows Col I fibrillation profile monitored by turbidity assay at pH 5 & 7, temperature 25 & 27 °C, & 0.05-0.1 mg/ml concentrations of Col I. (C) Temperature sensitive CD shows lack of proper Col I fibrillation due to pH 5 & temperature 25 °C at 0.1 mg/ml & lower concentration. (Di-ii) Representative thioflavin T (ThT) intensity plots show 100 µM Amylin aggregation has not been induced by presence of Col I (at 0.05 & 0.1 mg/ml conc.) at pH 5, temperature 25 °C & 37 °C.

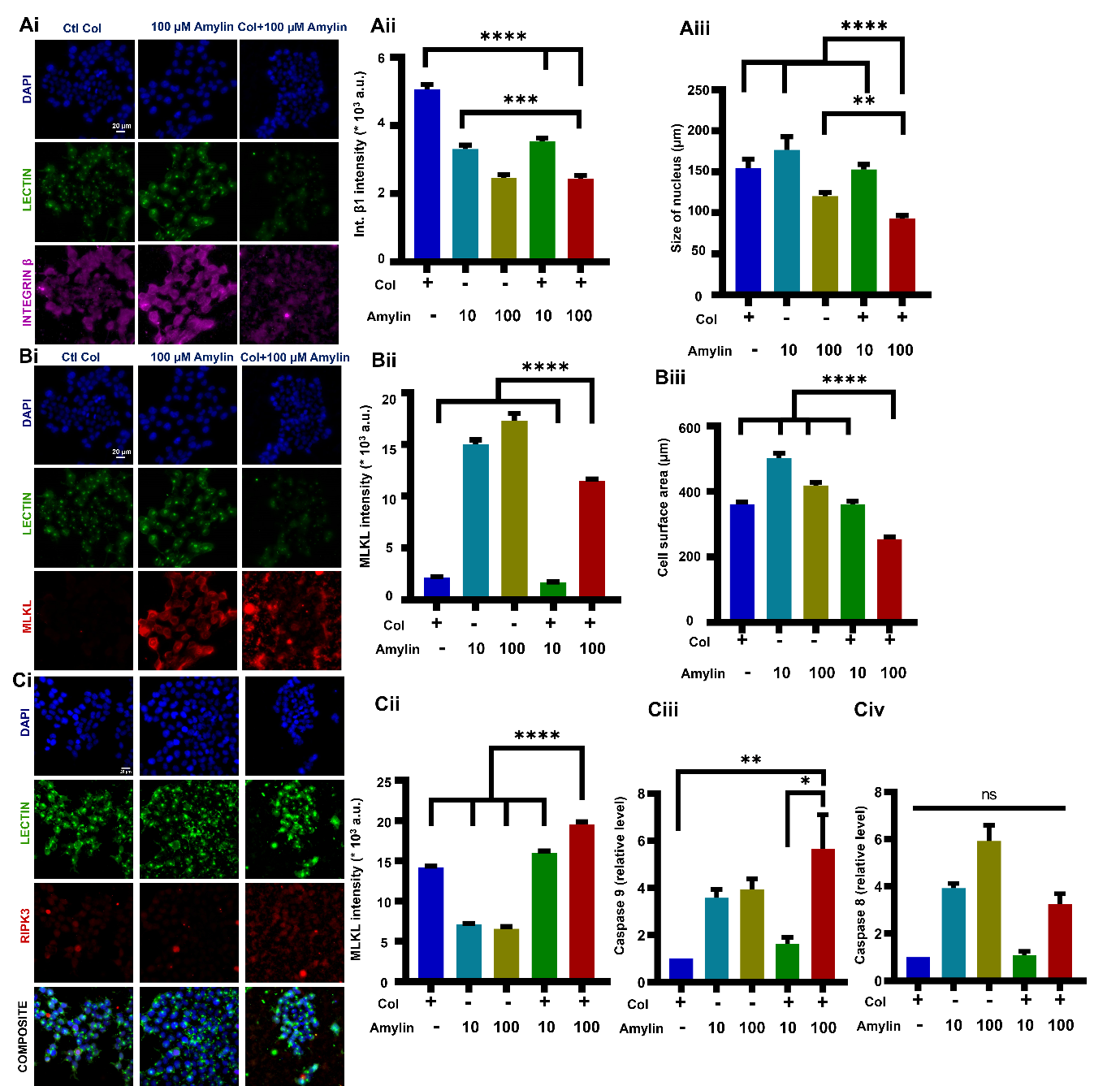

**Supp. Figure 4. Amyloid entangled matrices (AEMs) containing high amylin concentration are toxic to INS1E β cells.** (A) (i-ii) Representative integrin β1 (pink), lectin (green) and DAPI (blue) stained images of INS1E β cell across different conditions, and quantification of mean integrin β1 intensity ($\approx80-100$ cells per condition, $N=3$). (iii) Bar plot shows variability in the nucleus size across different conditions among which AEM100 possessing the lowest one. (B) (i-ii) Representative MLKL (red), lectin (green) and DAPI (blue) stained images of INS1E β cell across different conditions, and quantification of mean MLKL intensity ($\approx80-100$ cells per condition, $N=3$). (iii) Bar plot shows variability in the cell surface area across different conditions among which AEM100 possessing the lowest bar. (C) (i-ii) Representative RIPK3 (red), lectin (green) and DAPI (blue) stained images of INS1E β cell across different conditions, and quantification of mean RIPK3 intensity ($\approx80-100$ cells per condition, $N=3$). (iii-iv) Caspase 9 & caspase 8 genes expression in INS1E β cells across different conditions normalized to that on Col gels ($N=3$ per condition).

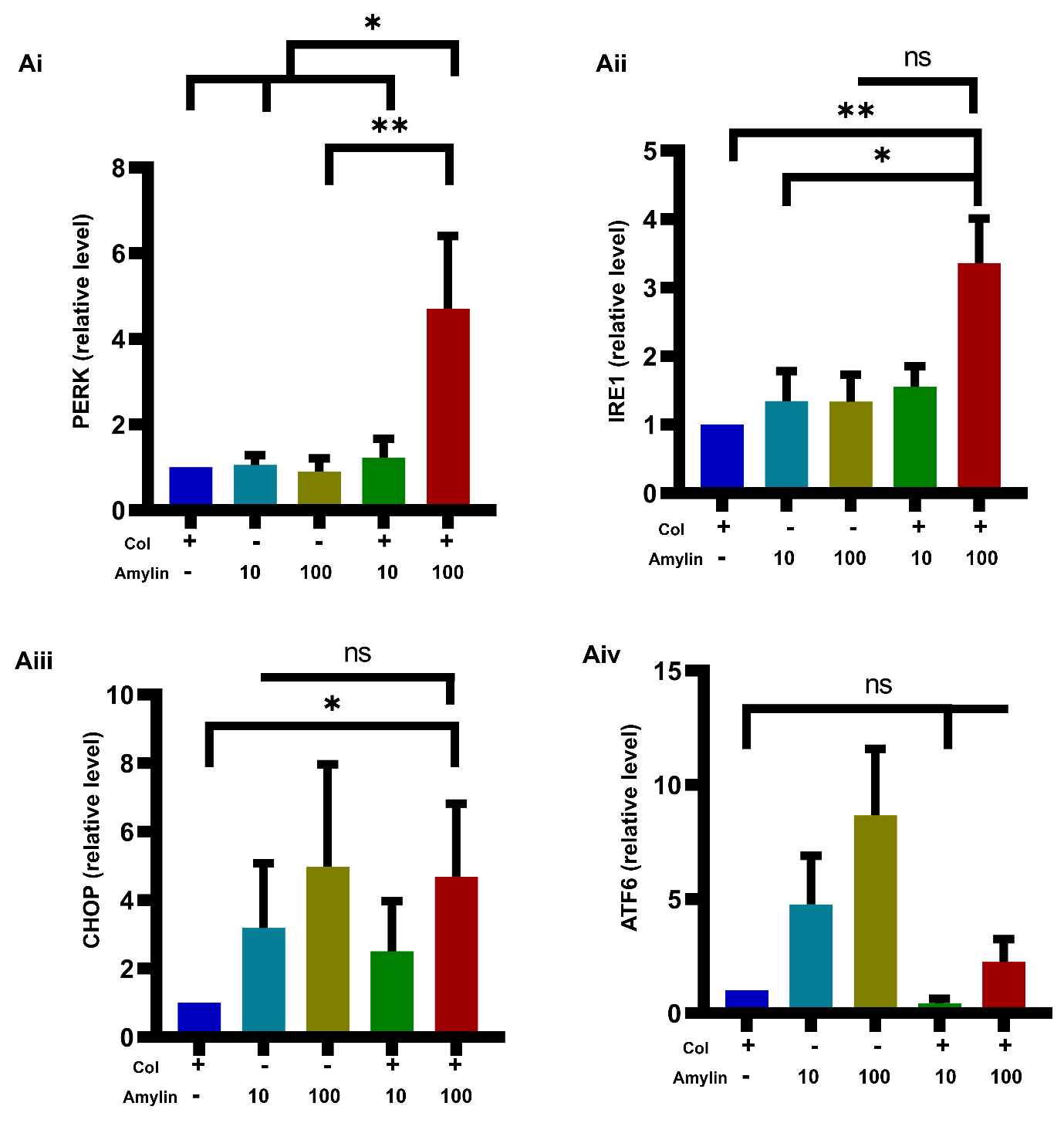

**Supp. Figure 5. Amyloid entangled matrices (AEMs) containing high amylin concentration causes endoplasmic reticulum (ER) stress to INS1E β cells.** (A) PERK, IRE1, CHOP, ATF6 genes expression in INS1E β cells across different conditions normalized to that on Col gels (*N* = 3 per condition).
